# Integrator dynamics in the cortico-basal ganglia loop underlie flexible motor timing

**DOI:** 10.1101/2024.06.29.601348

**Authors:** Zidan Yang, Miho Inagaki, Charles R. Gerfen, Lorenzo Fontolan, Hidehiko K. Inagaki

**Affiliations:** Max Planck Florida Institute for Neuroscience, Jupiter, FL 33458, USA; National Institute of Mental Health, Bethesda, MD 20814, USA; 3Turing Centre for Living Systems, Aix-Marseille University, INSERM, INMED U1249, Marseille, France

## Abstract

Flexible control of motor timing is crucial for behavior. Before volitional movement begins, the frontal cortex and striatum exhibit ramping spiking activity, with variable ramp slopes anticipating movement onsets. This activity in the cortico-basal ganglia loop may function as an adjustable ‘timer,’ triggering actions at the desired timing. However, because the frontal cortex and striatum share similar ramping dynamics and are both necessary for timing behaviors, distinguishing their individual roles in this timer function remains challenging. To address this, we conducted perturbation experiments combined with multi-regional electrophysiology in mice performing a flexible lick-timing task. Following transient silencing of the frontal cortex, cortical and striatal activity swiftly returned to pre-silencing levels and resumed ramping, leading to a shift in lick timing close to the silencing duration. Conversely, briefly inhibiting the striatum caused a gradual decrease in ramping activity in both regions, with ramping resuming from post-inhibition levels, shifting lick timing beyond the inhibition duration. Thus, inhibiting the frontal cortex and striatum effectively paused and rewound the timer, respectively. These findings suggest the striatum is a part of the network that temporally integrates input from the frontal cortex and generates ramping activity that regulates motor timing.

## Introduction

Flexible control of movement onset, or motor timing, is crucial for a wide range of behaviors, including vocal communication, driving, playing sports, and performing music. Most vertebrate species can adjust their motor timing to obtain reward or avoid punishment^1–9^, implying the ancient origins and significance of flexible motor timing. Without these abilities, our behaviors would be restricted to immediate reactions to stimuli.

To execute timed actions, the brain tracks the passage of time over seconds and then triggers actions, akin to how a timer beeps at the end of a preset duration. To perform actions at flexible timings, this ‘timer’ in the brain should be adjustable to the desired durations^4–10^. Neurons in the frontal cortex and striatum exhibit neural correlates of such a flexible timer. Before voluntary movements begin, many neurons in these areas demonstrate a gradual change in spiking activity, often characterized by ramping activity that peaks at the onset of movement^4–7,10–19^. When animals perform actions at various timings, the slope of this ramping activity varies accordingly, leading to the attainment of a hypothetical threshold level that triggers action at different timings^4–7,10–19^. Hence, the slow spiking dynamics in the frontal cortex and striatum may function as a timer, and the alteration in the speed of these dynamics could be analogous to adjusting the timer (Fig. 1a).

**Figure 1.**
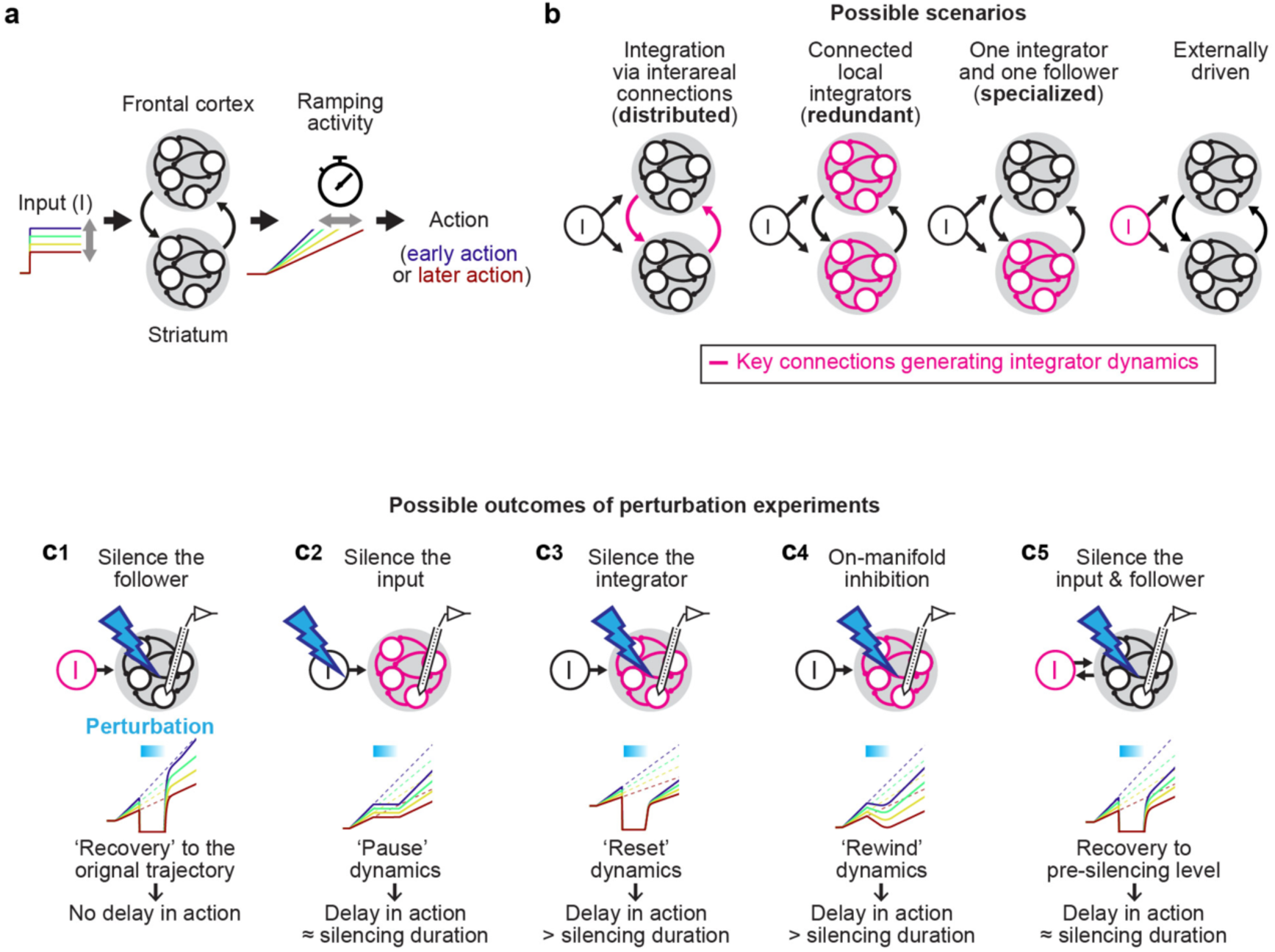
Multi-regional models of flexible motor timing. **a.** Schema of the computation performed by the cortico-basal ganglia loop for motor timing. The network may integrate a non-ramping (e.g., step) input to generate ramping dynamics with varying slopes, resulting in different lick timings. For simplicity, other areas connecting the striatum and frontal cortex are omitted. **b.** Possible configurations of the cortico-striatal network to implement integrator dynamics. The key connections that generate integrator dynamics are shown in pink. **c.** Schema of perturbation experiments and expected results.

Because isolated neurons can sustain activity only for tens of milliseconds, the seconds-long slow dynamics underlying motor timing likely arise from interactions among populations of neurons^19,20^. From a dynamical systems perspective, the population activity traces a trajectory within a high-dimensional state space, with each dimension corresponding to the activity of individual neurons^21^. Network interactions, for example via feedback connections, govern the evolution of trajectories and can stabilize certain states, known as attractors^19,20,22^. Within this framework, the slow dynamics can be generated by groups of quasi-stable states, forming a continuous attractor or point attractors with shallow basins. Such slow dynamics allow for the temporal integration of inputs to the network (when the input has a component aligned with the attractor), enabling the network to operate as an ‘integrator’^22–28^. Integrator networks can generate ramping activity by temporally integrating a non-ramping input (step or other types of inputs)^16,24,29^, with the amplitude of the input adjusting the speed of the ramp (Fig. 1a). Therefore, the integrator has been proposed as a mechanism to generate ramping activity for flexible motor timing^16,24,29^.

While neural correlates of an integrator have been observed in the frontal cortex and striatum^4,10,16,30^— two central brain areas in the cortico-basal ganglia loop — the precise roles of these areas in implementing the integrator dynamics remain unclear. Manipulations of the frontal cortex and striatum affect timing behavior, supporting their causal roles^10,18,31–41^. However, the presence of neural correlates and necessity cannot determine their computational roles^42,43^. First, neural correlates may be generated locally (internally generated) or inherited from other brain regions (externally driven). Second, a brain area that is ‘causal’ for timing behavior might 1) house the attractor landscape that generates integrator dynamics, 2) supply essential inputs for another brain area to function as an integrator, or 3) affect behavior through mechanisms independent of integrator dynamics, for example by controlling movement execution. This presents a general challenge in pinpointing the precise loci of computations within a multi-regional recurrent network. In the context of motor timing, the integrator generating the neural representation of time can either be: distributed across the frontal cortex and striatum; redundantly present in both areas; present in one area (specialized); or located in upstream areas (dynamics in the frontal cortex and striatum externally-driven) (Fig. 1b).

We addressed these knowledge gaps by performing a series of transient perturbations with simultaneous multi-regional electrophysiology. Depending on the computational role of the manipulated brain area, multi-regional dynamics are expected to respond to and recover from brief disturbances differently (Fig. 1c and Extended Data Fig. 1). For instance, silencing a brain area that is dispensable for the observed dynamics, i.e., an area with externally-driven dynamics, will result in a rapid return of ramping dynamics to the original trajectory after the silencing (Fig. 1c1). This manipulation has no effect on subsequent dynamics and actions, regardless of the inhibition’s strength, as observed during frontal cortical silencing in other behavioral tasks^44,45^. In contrast, inhibiting a brain area that is indispensable for the dynamics, such as one acting as an integrator or providing essential input, will result in a long-lasting change in subsequent dynamics and motor timing. The nature of this enduring influence can vary depending on the specific role of the targeted areas. Silencing a brain area that supplies essential input for an integrator will temporarily pause the integration in the recipient area, delaying the action by the duration of the silencing (Fig. 1c2). Silencing a brain area that acts as an integrator may reset the ramping dynamics, causing a delay in motor timing beyond the silencing duration (Fig. 1c3). In a specific situation, inhibition may target a combination of neurons that act as an integrator or input, altering activity patterns along the direction in the activity space implementing the integrator. This ‘on-manifold’ inhibition^43^ will be integrated into the ramping dynamics, resulting in the rewinding of the representation of time during the inhibition and causing a delay in motor timing beyond the silencing duration (Fig. 1c4). Additionally, a brain area can function as both the input and follower of an integrator (‘input/follower’). In this case, silencing the input/follower area will pause integration in the connected integrator. Thus, when the input/follower area recovers, its activity will return to the pre-silencing level (but not the original trajectory), causing a delay in action equal to the duration of the silencing (Fig. 1c5).

To systematically dissect multi-regional dynamics following this model-driven approach, we developed a flexible lick-timing task in mice, where mice explore various lick times over 600 trials per session (652 ± 9 trials; mean ± SEM; 224 sessions, 48 mice; Fig. 2a). This enabled a series of perturbations within individual sessions. Large-scale electrophysiology in the frontal cortex and striatum allowed decoding of planned lick time in individual trials, providing an ideal testbed to quantify how the dynamics respond to perturbations. Leveraging this system, we identified specialized functional roles of the frontal cortex and striatum in implementing integrator dynamics, generating ramping activity that serves as an adjustable timer.

**Figure 2.**
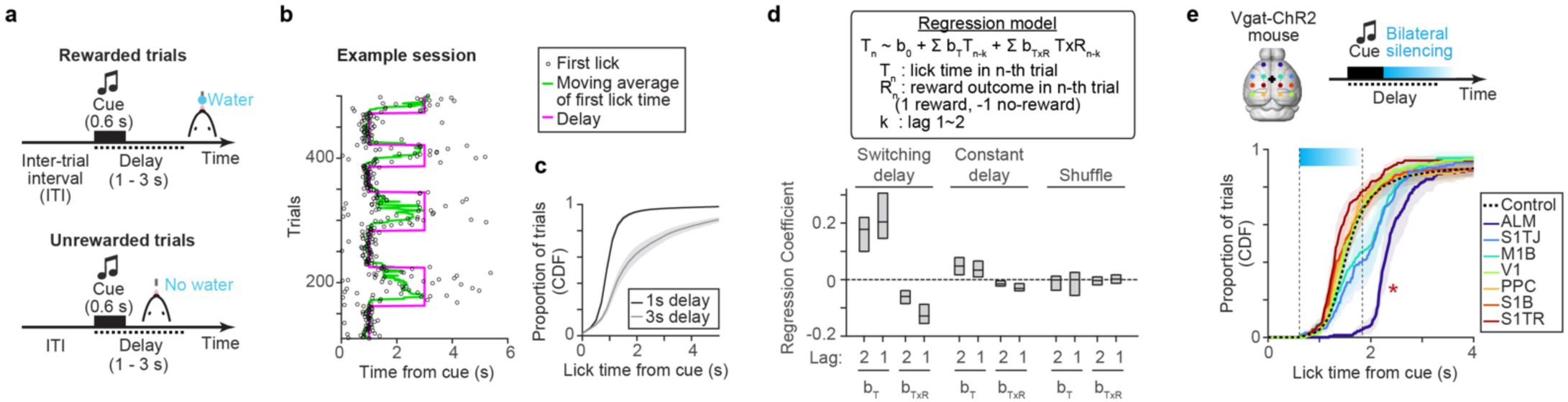
ALM is required for lick timing control. **a.** Flexible lick timing task. The delay epoch started at the trial onset signaled by the cue. The first lick after the delay was rewarded, and a premature lick during the delay aborted the trial. **b.** Example session. Only part of a session is shown for visualization. **c.** Cumulative distribution of lick time in 1 s and 3 s delay blocks (56 sessions, 10 mice). Shades, SEM (hierarchical bootstrap). **d.** Regression coefficients of the regression model based on previous lick time (T) and its interaction with reward outcome (T × R), with 2-trial lags. Switching delay (n = 30 mice). Constant delay (n = 13 mice). The central line in the box plot, median. Top and bottom edges, 75% and 25% points. **e.** Top, optogenetic loss-of-function screening of dorsal cortical areas during delay. Bottom, cumulative distribution of lick time in trials with different silenced areas (3879 control trials, 172 ± 5 silencing trials per brain region; mean ± SEM; 9 sessions, 3 mice). Cyan bar, silencing window (1.2 s). Shades, SEM (hierarchical bootstrap). *, *p* < 0.001 (hierarchical bootstrap with a null hypothesis that the lick times in control trials are later than or equal to the ones in silencing trials). Regions adjacent to ALM (M1B and S1TJ) exhibited weaker effects, attributed to the limited spatial resolution of the manipulation^48^.

## Results

### The frontal cortex is required for lick timing control

We developed a flexible lick-timing task for mice. In this task, an auditory cue (3 kHz, 0.6 s) signals trial onset followed by a delay epoch with unsignaled duration (1 - 3 s; Fig. 2a, Methods). If mice lick after the delay, they receive a water reward. In contrast, a premature lick during the delay terminates the trial without reward. We varied the delay duration in blocks of trials within a session (with the number of trials within each block randomly selected from 30 - 70 trials; Fig. 2b and Extended Data Fig. 2ab). Despite the absence of a sensory cue instructing the delay duration or block transitions, mice dynamically adjusted their lick time distribution according to the delay within 10 trials after the delay switch (Fig. 2bc and Extended Data Fig. 2de).

The only information available for mice to infer the appropriate lick time in this task is their prior lick times and the outcomes. To investigate whether such ‘trial history’ guides trial-by-trial lick timing, we exploited a linear regression model to predict future lick times based on trial history^31,46^ (n = 30 mice; Methods). This analysis revealed a positive correlation between upcoming lick times and previous lick times, alongside a negative influence of previous reward outcomes on upcoming lick times: mice tended to lick earlier after a reward and later after a lack of reward (Fig. 2d and Extended Data Fig. 2l). Since an absence of reward indicates a premature lick in this task, delaying licking after an unrewarded attempt is an adaptive strategy. In contrast, when we keep the delay duration constant across trials and sessions (‘constant delay condition’; n= 13 mice; Extended Data Fig. 2c), there was no significant influence of former trials on lick timing (Fig. 2d). Thus, mice use trial history to strategically adjust lick timings^32^ only when the delay duration is variable. We employed this behavior as a model system to examine how the brain strategically and flexibly adjusts the timing of actions (see Extended Data Figs. 2 and 3 for further quantifications, models explaining lick time distribution, and body part tracking).

To identify dorsal cortical areas that control lick timing in an unbiased manner, we performed optogenetic loss-of-function screening using transgenic mice expressing ChR2 in GABAergic neurons (Vgat-ChR2-EYFP mice^47^) with a clear skull preparation^48^ (Methods). We bilaterally silenced individual dorsal cortical areas during the delay epoch (delay duration: 1.5 s) by scanning a blue laser in randomly interleaved trials (488 nm, 1.5 mW). Photostimulation started 0.6 s after the cue and lasted for 1.2 s including a 0.2 s ramp down of laser power at the end (3 mice; Fig. 2e; Methods). Lick initiation was blocked during the silencing of a frontal cortical area: anterior-lateral motor cortex (ALM; anterior 2.5 mm, lateral 1.5 mm from Bregma). This is consistent with the established role of ALM as a premotor cortex for licking^19^. Notably, following the transient silencing of ALM, the distribution of lick times shifted significantly later, with the median lick time delayed by 0.79 (0.59 - 0.97) s (mean and 95% confidence interval). This suggests a causal role of ALM in controlling the lick time.

### Similar scalable timing dynamics in ALM and striatum

To investigate neural activity associated with lick time control in ALM, we conducted high-density silicon probe recordings (4467 putative pyramidal neurons recorded in 172 sessions, 45 mice; Extended Data Table. 1). Many ALM neurons displayed ramping activity during the task, peaking around the onset of lick (Fig. 3a; Out of 4467 neurons, 1363 showed a significant increase and 1809 showed a significant decrease in spike rate before the lick compared to the baseline; Methods). The ramping speed of these neurons varied across trials and predicted lick timing (Fig. 3a). Notably, temporal warping^10,16,36,41,49–52^, which normalizes the temporal axis between cue and lick in each trial, significantly reduced across-trial variability in spike rate in 68.1% of neurons (Fig. 3a bottom and Extended Data Fig. 4a-c). This indicates that two-thirds of ALM neurons exhibited temporal scaling (stretching or shrinking) of activity patterns across trials, with the speed of their dynamics anticipating lick time.

**Figure 3.**
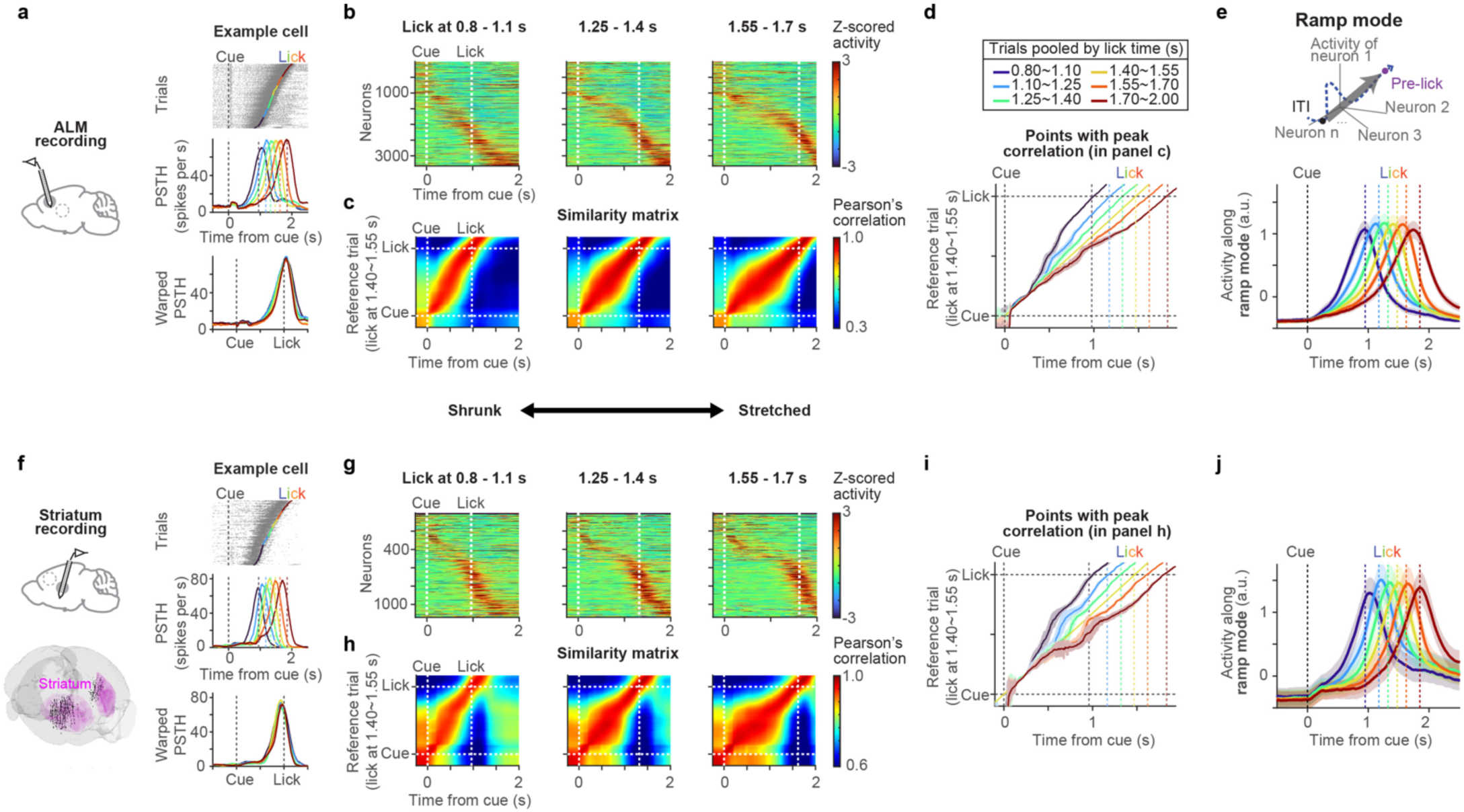
Similar scalable timing dynamics in ALM and striatum. **a.** An example ALM cell. Top, spike raster. Trials are sorted by lick time. Colored dots, first lick time, divided into six lick time ranges shown in **d** (top). Middle, peri-stimulus time histogram (PSTH) of the six lick time groups. Vertical dotted lines in the same color indicate the corresponding lick time. Bottom, PSTH following temporal warping (Methods). **b.** ALM activity (Z-scored) across trials with different lick time ranges. Neurons are sorted in the same order for all panels (by peak firing time in trials with lick between 1.40 - 1.55 s). Only neurons with more than 10 trials across all six lick time ranges were analyzed (n = 3261 neurons, 45 mice). **c.** Similarity matrix (Pearson’s correlation) comparing ALM population activity between trials with different lick time ranges. **d.** Time points with the peak correlation in the similarity matrix (**c**), across six different lick time ranges. Lines, mean. Shades, SEM (hierarchical bootstrap). **e.** Top, schema of ramp mode. Bottom, ALM population activity projected along the ramp mode, with trials grouped by lick time (same color scheme as **d**). n = 3261 neurons, 45 mice. Lines, grand mean. Shades, SEM (hierarchical bootstrap). **f-j.** The same as in **a**-**e** but for striatal recording (n = 1073 neurons, 16 mice). Bottom in **f**, the location of all recorded units registered to Allen common coordinate framework.

Consistent with individual cells, the population activity patterns in ALM appeared to temporally scale with lick time (Fig. 3b). To quantify this observation, we calculated similarity matrices using Pearson’s correlations, comparing population activity between trials with differing lick times (Fig. 3c). We then identified the points with peak correlations in these similarity matrices (Fig. 3d). This analysis revealed that although similar population activity patterns emerged across trials, the speed at which these patterns unfolded varied as a function of lick time (Fig. 3d).

To further characterize the population activity patterns between the cue and the lick, we defined three modes (directions in population activity space) to differentiate the spike rate during specific time windows after the cue from the baseline activity during the inter-trial interval (ITI; 0 - 1 s before the cue): 0 - 300 ms after the cue (cue mode, CM), 500 - 800 ms before the lick (middle mode, MM), and 200 - 500 ms before the lick (ramp mode, RM) (Methods). Population activity along these three modes collectively spanned the time from cue to lick and explained most of the task-modulated activity (74%; Extended Data Fig. 5). Notably, population activity along the MM (MM activity) and RM (RM activity) displayed temporal scaling (Fig. 3e and Extended Data Fig. 5b-d, k-l). Thus, a large proportion of ALM activity between the cue and the lick, which we refer to as ‘timing dynamics’, demonstrated temporal scaling, similar to observations in other timing tasks and species^10,16,36,41,49–52^.

Many excitatory neurons in ALM project to the striatum, spanning various sectors, including the ventrolateral striatum^53–55^ (VLS; Extended Data Fig. 6a). Considering the involvement of the striatum in various timing behaviors^10,13,18,33,34,36,38–40,49,50^, it likely cooperates with ALM to control lick timing. We performed electrophysiological recordings in the striatum using Neuropixels probes^56^ (Fig. 3f and Extended Data Fig. 6b and d). In total, we recorded 1972 neurons (97 sessions, 16 mice), with the majority classified as putative striatal projection neurons (SPN; 64%) or fast-spiking interneurons (FSI; 30%) based on spike features^57^ (Extended Data Fig. 6e). Because both SPN and FSI showed similar activity patterns (consistent with previous study^57^) and comprised ∼95% of the population together, we have pooled all cell types for analysis. Overall, striatal activity between the cue and lick was analogous to those in ALM^10,18,49,57^. Firstly, the activity of individual striatal neurons was well-explained by temporal scaling (in 56% of cells; Fig. 3fg and Extended Data Fig. 4c), and many neurons displayed ramping activity (out of 1972 neurons, 741 showed a significant increase and 400 showed a significant decrease in spike rate before the lick compared to the baseline). Secondly, striatal population activity (similarity matrix, MM activity, and RM activity) showed a temporal profile similar to that in ALM with temporal scaling (Fig. 3h-j and Extended Data Fig. 5f-l).

In some sessions, we simultaneously recorded from ALM and the striatum (16 sessions, 8 mice). At each time point, we used the population activity from either ALM or the striatum to predict how much time was left until lick using a k-nearest neighbor (kNN) decoder (‘time to lick (T_to lick_)’; Extended Data Fig. 6f-g; Methods). The decoded T_to lick_ in these two brain areas was significantly correlated across individual trials (Extended Data Fig. 6h-j). Thus, ALM and the striatum demonstrate similar scalable timing dynamics coupled at the single-trial level.

### Neural correlates of trial history in ALM and striatum

What determines the speed of timing dynamics after the cue and guides lick time? Given that trial history influences lick time in this task (Fig. 2d), we hypothesized that certain neurons encode trial history before the cue. Such neural activity could establish the initial conditions of a network^9,46^ and/or provide inputs^31^ to guide the speed of timing dynamics and action timing.

In line with our hypothesis, some ALM neurons exhibited tonic activity during ITI, predicting the upcoming lick time even 2 - 3 s before the cue onset: the proportion of neurons with a significant rank correlation of spiking activity during ITI vs. upcoming lick time was 22.8 (19.4 - 26.5)%; mean (95% confidence interval; Extend Data Fig. 4e-g; Methods). The tonic activity of these neurons was correlated with the lick time and reward outcome of the previous trial (Fig. 4a and Extend Data Fig. 4ef). Because upcoming lick time and trial history are correlated (Fig. 2d), we calculated the partial correlation between the activity of ALM neurons and previous lick time while removing the effect of upcoming lick time (Methods). This partial correlation was significantly higher than trial shuffle and session permutation controls^58^ both before and after the cue (Extended Data Fig. 4h). ALM neurons encoding previous lick time also tended to encode previous reward outcome (Extended Data Fig. 4i). Together, ALM neurons encode trial history and anticipate upcoming lick time even before the cue.

**Figure 4.**
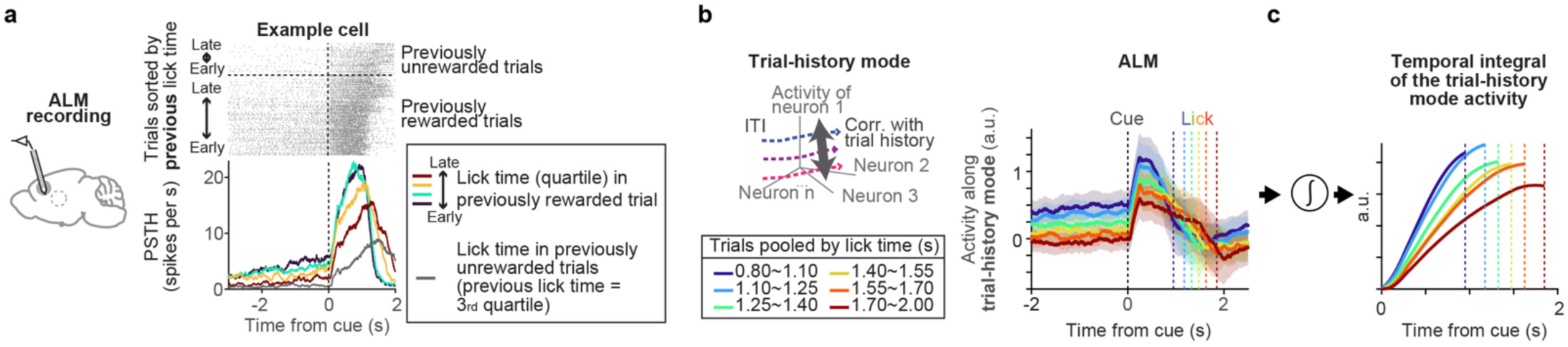
Neural correlates of trial history. **a.** An ALM example cell whose activity is modulated by the lick time and reward outcome in the previous trial. Top, spike raster, grouped by reward outcome in the previous trial and sorted by the lick time in the previous trial. In this example cell, the ITI activity is higher in trials after rewarded trials and with earlier licks. Bottom, PSTH. Lick times of the previous rewarded trials were divided into quartiles indicated by different colors. The gray trace, trials following previously unrewarded trials with previous trial’s lick times within the 3rd quartile. **b.** Left, schema of trial-history mode. Right, ALM population activity projected along trial-history mode. n = 3261 neurons, 45 mice. Lines, grand mean. Shades, SEM (hierarchical bootstrap). **c.** Temporal integration of activity along the trial-history mode after the cue (based on plots in **b**) results in ramps with different slopes.

If trial-history information encoded in ALM influences subsequent lick times, such neural correlates may be absent in contexts where trial history is not used. Supporting this idea, ALM activity was not correlated with trial history above chance level under the constant delay condition (Extended Data Fig. 4h and l), where mice did not rely on trial history to guide their lick times.

To examine the evolution of ALM activity encoding trial history at the population level, we defined a “trial-history mode” defined by the rank correlation between the ITI activity and trial history (the predicted lick time based on previous trials using trial-history regression; Fig. 4b, left; Methods). ALM activity along the trial-history mode was modulated after reward outcome (Extended Data Fig. 4j_3_), exhibited a graded persistent activity during the ITI, and showed a step-like increase upon the cue presentation at the trial onset (Fig. 4b). The amplitude of this activity anticipated upcoming lick times throughout the trial (Fig. 4b). Thus, activity along this mode carries trial-history information and predicts upcoming lick time across trials. Temporal integration of graded activity with varying amplitudes produces a ramp with different slopes (with the cue at trial onset acting as a gate to initiate integration). Indeed, temporal integration of the activity profile of the trial-history mode after the cue generates ramping with different slopes (Fig. 4c). Similarly, some striatum neurons also exhibited tonic activity during ITI anticipating upcoming lick time (but it was weaker than in ALM; Extended Data Fig. 4g and k). Together, there are neural correlates of trial history in both ALM and striatum, which may function as an input to the integrator and guide the speed of timing dynamics and lick timing.

### ALM silencing pauses the timer

Neural correlates of temporal integration (i.e., trial-history mode and RM) in ALM alone are insufficient to conclude that ALM is the integrator. Likewise, while ALM silencing shifted lick time (Fig. 2e), this behavioral effect alone is insufficient to attribute a specific computational function to ALM (Extended Data Fig. 1). Therefore, to examine whether ALM functions as an integrator, we recorded ALM activity using silicon probes during calibrated silencing.

Strong cortical silencing can induce post-silencing rebound spiking activity that triggers actions^59,60^ complicating the interpretation of subsequent dynamics and behavior. To address this, we calibrated the silencing protocol to minimize rebound activity. Silicon probe recordings confirmed nearly complete ALM silencing during photostimulation using 1.5 mW 488 nm laser in Vgat-ChR2-EYFP mice (8 spots, bilateral). Limiting the silencing duration to 0.6 s (including a 0.3 s ramp down) minimized post-silencing rebound activity and licking (Extended Data Fig. 7a-d). However, rebound activity and licking increased over sessions due to an unknown adaptation mechanism^61,62^ (Extended Data Fig. 7ab). Therefore, we restricted our analysis to the initial two days of ALM silencing (similarly, restricted to the first day for striatal manipulation in Fig. 6; Extended Data Fig. 7).

Silencing ALM during the delay epoch (0.6 s after cue onset) with this protocol shifted the median lick time by 0.47 (0.37 - 0.56) s (mean and 95% confidence interval, n = 14 mice; Fig. 5ab and Extended Data Fig. 8a-d). Interestingly, this temporal shift was close to the silencing duration (0.6 s), as if the ‘timer’ was paused during ALM silencing. In contrast, silencing ALM before the cue did not shift the lick time distribution, suggesting that this manipulation has no long-lasting effect on lick time (ALM activity rapidly recovered after ITI silencing, implying that trial-history information is robustly maintained across brain areas before the trial onset^63–66^; Extended Data Fig. 9q).

**Figure 5.**
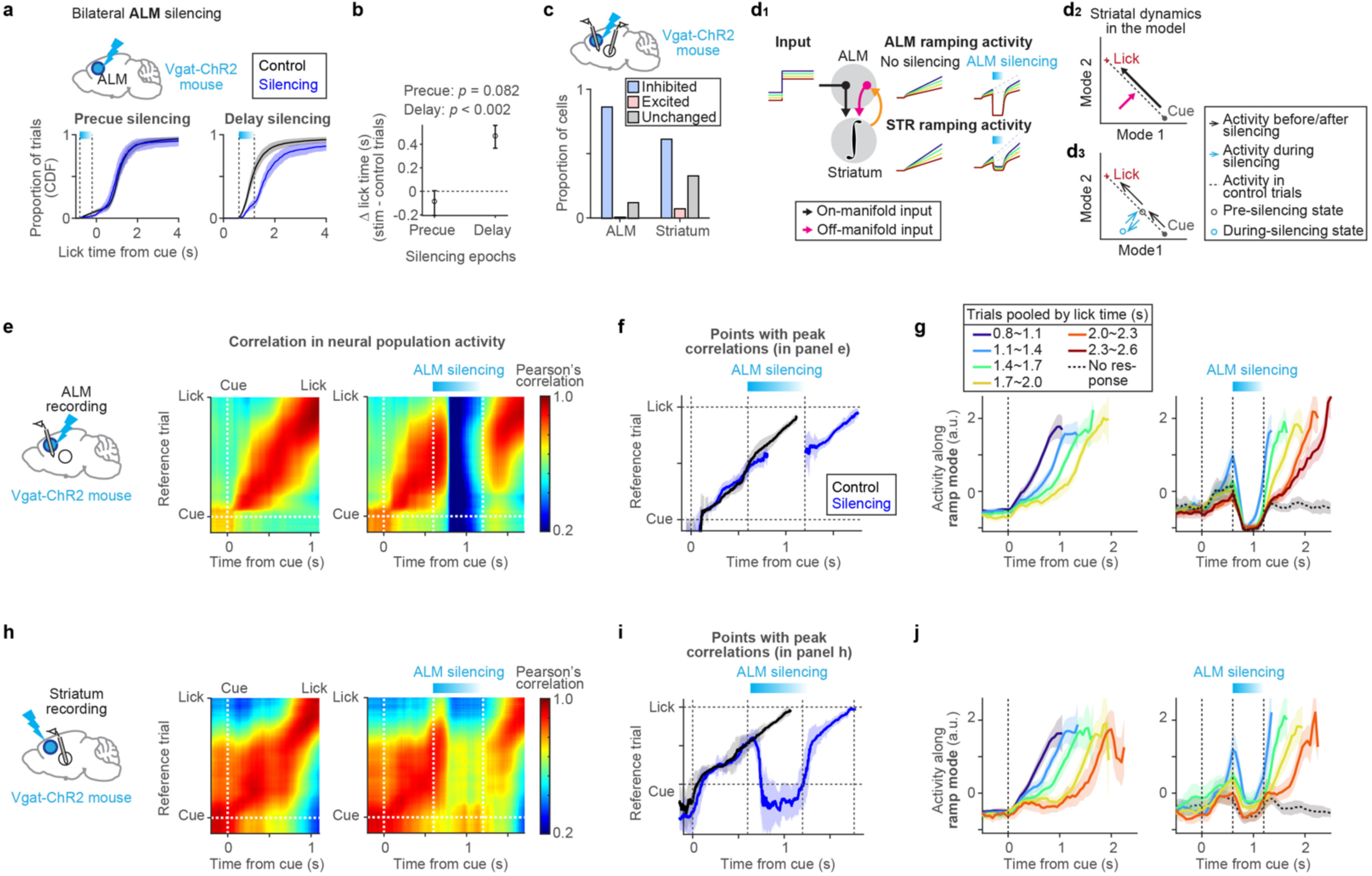
Transient optogenetic perturbation of ALM. **a.** Top, bilateral ALM silencing in Vgat-ChR2 mouse. Bottom, cumulative distribution of lick time in trials with precue (left) or delay silencing (right). Black, control trials. Blue, silencing trials. Shades, 95% confidence interval (hierarchical bootstrap). N = 28 sessions, 14 mice. **b.** Change in median lick time in **a**. Error bars, 95% confidence interval (hierarchical bootstrap). *P*-value, hierarchical bootstrap with a null hypothesis that there is no change in lick time. **c.** The proportion of ALM pyramidal neurons (left) and striatal neurons (right) with spiking activity significantly inhibited or excited (*p* < 0.05; Unchanged, *p* >= 0.05; rank sum test) during ALM silencing. Neurons with a mean spike rate above 1 Hz in control trials during the silencing were analyzed. This includes 391 neurons in ALM (n = 28 sessions, 14 mice) and 238 neurons in the striatum (n = 14 sessions, 7 mice). **d.** Schema of a model explaining the data (**d1**). Schema of how on- and off-manifold inputs from ALM modulate activity in striatum (**d2**). Schema of striatal activity during ALM silencing (**d3**). Orange arrows, ALM receives input from the striatum to follow the ramping activity generated there. **e.** Left, recording of ALM during ALM silencing. Middle and right, similarity matrix of ALM population activity between trial types. Middle, control (unperturbed trials with lick around the median lick time; Methods) vs. ref trial (unperturbed trials with lick between 1.4 - 1.7 s). Right, ALM silencing trials (perturbed trials with lick between 1.7 - 2.0 s) vs. ref trial. **f.** Points with the peak correlation in the similarity matrix in **e**. Lines, mean. Shades, SEM (hierarchical bootstrap). Black, control. Blue, silencing. **g.** ALM population activity along the ramp mode (n = 28 sessions, 14 mice), trials grouped by lick time. Activity up to lick time is shown. Lick time ranges that exist in at least two-thirds of the analyzed sessions were shown. Therefore, the plotted lick time ranges vary depending on the manipulation conditions. Lines, grand mean. Shades, SEM (hierarchical bootstrap). Left, control. Right, ALM silencing. **h-j.** Same as in **e-g** but for striatal recording. n = 14 sessions, 7 mice.

Silicon probe recordings of ALM during delay silencing (590 neurons, 14 mice) revealed that, despite near-complete silencing during photostimulation, population activity patterns resembling those just before the silencing suddenly reemerged after the silencing ceased, and ALM dynamics continued to unfold from there in parallel to the unperturbed condition (quantified by similarity matrix; Fig. 5ef). Consistently, at the end of ALM silencing, RM activity rapidly recovered (Fig. 5g and Extended Data Fig. 9a). However, RM activity after silencing was significantly lower than in the unperturbed condition (Extended Data Fig. 9a), indicating that it is not a recovery to the original trajectory as depicted in Fig. 1c1. Instead, RM activity recovered close to the pre-silencing level, after which activity began to slowly ramp up again (Fig. 5g and Extended Data Fig. 9a; akin to Fig. 1c5). To appreciate high-dimensional timing dynamics, we also used a kNN decoder to analyze the evolution of decoded T_to lick_. We compared ALM silencing and unperturbed trials with matched decoded T_to lick_ before the silencing (0.6 s after the cue). During the silencing, the decoded T_to lick_ diverged between ALM silencing and unperturbed trials, and such shift persisted in parallel after the silencing (Extended Data Fig. 9d). Together, after ALM silencing, ALM activity rapidly recovered to the pre-silencing level, and then ALM dynamics unfolded in parallel to those observed under unperturbed conditions, explaining the shift in lick time close to the silencing duration.

The rank order of RM activity across trials predicted lick time under unperturbed conditions (Fig. 5g, left). Notably, this rank order collapsed during silencing but recovered afterward (Fig. 5g and Extended Data Fig. 9b). Similarly, ALM population activity before and after silencing both significantly predicted upcoming lick time on single trials using the kNN decoder, whereas this predictability was lost during silencing (Extended Data Fig. 9c; Methods). These findings suggest that following silencing, ALM activity rapidly reverted to a state closely resembling its pre-perturbation condition at the individual trial level.

The recovery of ALM activity to pre-silencing levels and the shift in lick time close to the silencing duration challenge multiple network models where ALM acts as the only integrator or is solely driven by external inputs (Extended Data Fig. 1). Instead, our data suggest that ALM silencing momentarily pauses the temporal integration because ALM provides input to an integrator (a ‘timer’) in another brain area. Additionally, the rapid post-perturbation recovery of ALM ramping activity to pre-perturbation levels implies that ALM ramping activity follows the dynamics of the external integrator, which was paused during the ALM silencing. This indicates that ALM acts both as an input to and a follower of the external integrator (Fig. 1c5; Extended Data Fig. 1f).

### The striatum maintains timing information during ALM silencing

During ALM silencing, other brain areas must retain the timing information to restore ALM dynamics afterward. Given the prominent timing dynamics observed in the striatum (Fig. 3), the striatum may serve such a role. To test this, we recorded striatal activity during ALM silencing using Neuropixels probes^56^ (372 neurons, 7 mice). A significant portion of striatal neurons (60%) decreased their spike rates during ALM silencing, indicating that ALM provides a major excitatory drive to the striatum (Fig. 5c, right, and Extended Data Fig. 10e-h). Consistently, upon ALM silencing, RM activity in the striatum rapidly decayed, and the population activity patterns (similarity matrix) resembled those observed during ITI when the spike rate was low (Fig. 5h-j). Following ALM silencing, striatal activity recovered to near pre-silencing levels, and then dynamics unfolded in parallel to those observed under unperturbed conditions (Fig. 5h-j), similar to what was observed in ALM. However, striatal activity was not entirely abolished during ALM silencing (Fig. 5c and j). The residual striatal activity during ALM silencing maintained the rank order of RM activity and predicted lick time (Fig. 5j and Extended Data Fig. 9fg). Thus, despite the reduced mean spiking activity, the striatum retained timing information during ALM silencing.

Striatum activity stayed at a low level and did not ramp up during ALM silencing (Fig. 5j), suggesting ALM is providing essential input to generate the ramping activity. The observed multi-regional dynamics can be replicated by a model where the striatum (and/or subcortical areas situated between the striatum and ALM, such as the substantia nigra reticulata and the thalamus) functions as an integrator, and ALM acts as both an input to and a follower of this ‘subcortical integrator’ (Fig. 5d1 and Extended Data Fig. 1f). In this model, input from ALM to the striatum has both on- and off-manifold components to explain the pause of time representation and the reduction in striatal activity during ALM silencing:

1. The on-manifold component of the ALM input is temporally integrated by the subcortical integrator to generate scalable timing dynamics (Fig. 5d_1-2_, black arrows). The trial-history mode activity may play this role (Fig. 4bc). In the state space, this input aligns with the direction in which timing dynamics evolve (i.e., along the direction of integration, Fig. 5d2). During ALM silencing, the lack of on-manifold input pauses the evolution of activity along this direction, effectively pausing the representation of time.
2. The off-manifold component of the ALM input functions as an excitatory drive (Fig. 5d_1-2_, pink arrows). In the state space, the off-manifold input is orthogonal to the direction in which timing dynamics evolve, amplifying striatal activity without affecting the representation of time (Fig. 5d2). During ALM silencing, the lack of this excitatory drive results in a large reduction in striatal activity.

Altogether, due to the loss of these inputs from ALM, the representation of time is paused in the striatum at a reduced activity level during ALM silencing (cyan circle in Fig. 5d3). Once ALM silencing ends, the excitatory drive returns, and the striatal activity recovers to pre-silencing levels. Additionally, the recovery of the on-manifold input after ALM silencing allows the timing dynamics to evolve from the pre-silencing level along a normal trajectory. This results in a parallel shift in timing dynamics after ALM silencing, consistent with the experimental data (Fig. 5d, and Extended Data Fig. 1f).

Temporal integration can be achieved through feedforward (FF) networks^67–69^, in addition to positive feedback loops (Fig. 5d and Extended Data Fig. 1). Recurrent network models with FF connections can produce both sequential evolutions of population activity patterns and ramping activity, as observed in the data (Extended Data Fig. 11a-c). We generated multi-regional FF networks assigning different roles to ALM and the striatum. In a model where ALM acts as an input/follower of the subcortical FF network, ALM perturbation resulted in a pause in time representation similar to the data (but not in other models; Extended Data Fig. 11d-g). In this model, ALM provides the recurrent excitatory input necessary for the activity patterns to unfold in the subcortical FF network. Without this input, the representation of time pauses. Therefore, regardless of the detailed implementation of the integrator (i.e., positive feedback models or FF models), our data suggest that ALM provides essential input for the subcortical integrator to maintain temporal integration and represent time. The key assumption of these models is that the striatum (and/or subcortical areas situated between the striatum and ALM) serves as the integrator. To test this prediction, we conducted perturbation experiments in the striatum.

### Striatum inhibition rewinds striatal timing dynamics

The striatum contains two major projection cell types: D1 receptor-expressing direct pathway SPN (D1-SPN) and D2 receptor-expressing indirect pathway SPN (D2-SPN)^70^. Consistent with the anti-kinetic function of D2-SPN^70–72^, D2-SPN silencing and non-cell-type-specific striatal silencing using the soma-targeted light-dependent chloride channel, stGtACR1^73^, triggered licking during photostimulation (Extended Data Fig. 8m-s). This response made it unsuitable for testing the function of the striatum in lick timing control. Consequently, we focused on inhibiting D1-SPNs to investigate the effects of transient striatal perturbation.

To inhibit D1-SPN, we crossed Drd1-cre FK150^74^ line with cre-dependent stGtACR1 transgenic reporter mice^59^, and bilaterally implanted tapered fiber optics^75^ (1.0 mm taper, NA 0.37) to deliver light to a large volume in the striatum (Fig. 6b and Extended Data Fig. 6c). We confirmed the inhibition using optrode recordings (488 nm, 0.25 - 0.5 mW): 9 out of 25 SPNs (36%) significantly reduced spike rates without axonal excitation^59,76,77^ or post-silencing rebound (Fig. 6e, left, Extended Data Fig. 7de, and Extended Data Fig. 10i-l). We cannot distinguish SPN subtypes based on spike features^55,78^, but since half of the SPNs are D1 SPN^70^, we estimated that we inhibited ∼70% of D1 SPN around the fiber optics.

**Figure 6.**
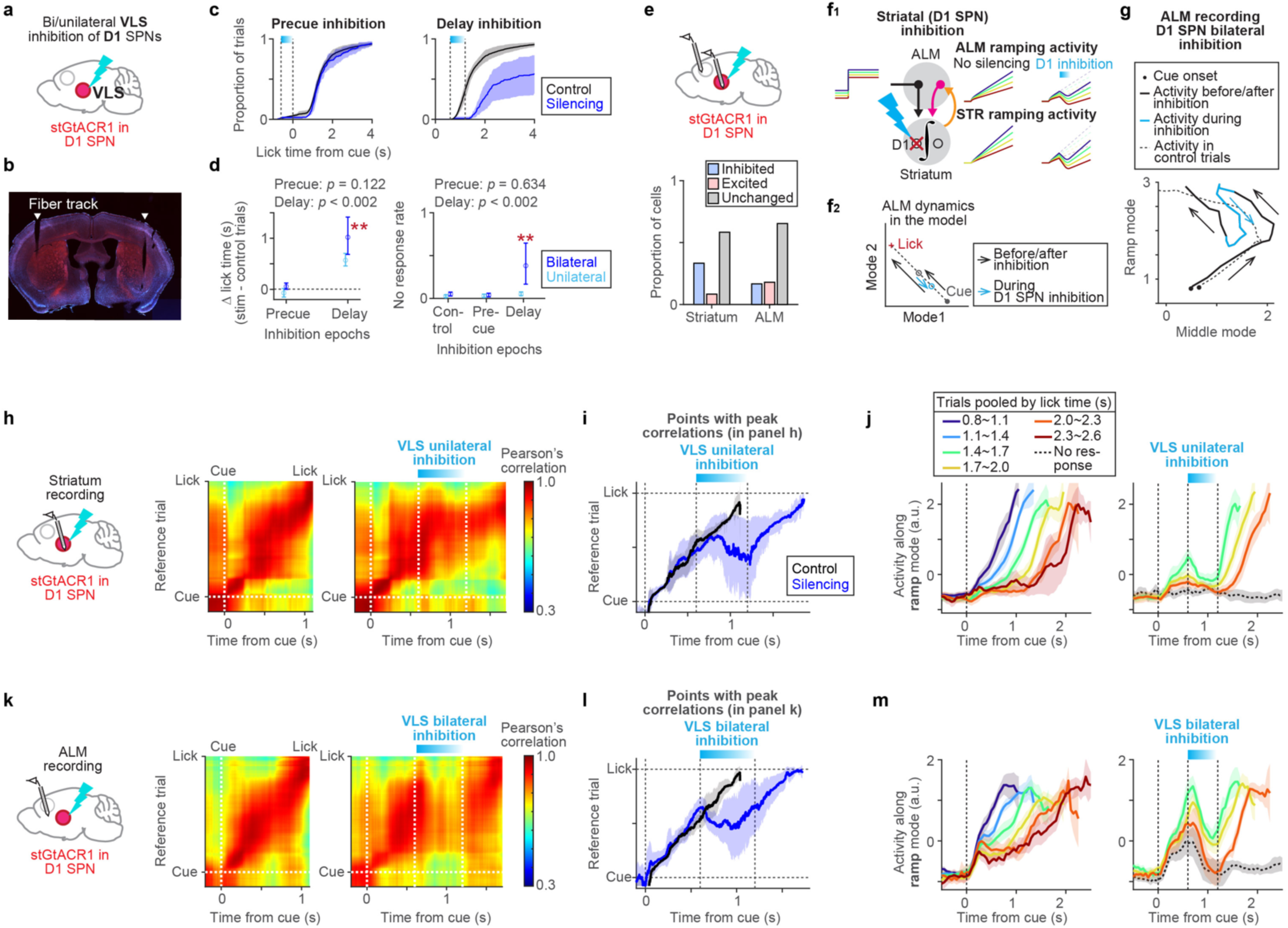
Transient optogenetic perturbation of D1 SPN in the striatum. **a.** Schema of inhibiting D1 SPN in ventral lateral striatum (VLS). **b.** Coronal section of a Drd1-cre;cre-dependent-stGtACR1:fusion-red mouse with bilateral tapered optic fiber implanted in the VLS (white arrows). Red, fusion red. Blue, DAPI staining. **c.** Cumulative distribution of lick time in trials with precue (left) or delay inhibition (right). Black, control trials. Blue, trials with bilateral inhibition. Shades, 95% confidence interval (hierarchical bootstrap). **d.** Left, change in median lick time in **c**. Right, no-response rate in **c**. Error bars, 95% confidence interval (hierarchical bootstrap). *P*-value, hierarchical bootstrap with a null hypothesis of no change compared to control in bilateral inhibition. **, *p* < 0.002. Blue, bilateral inhibition. Light blue, unilateral inhibition. **e.** The proportion of striatal neurons (left) and ALM pyramidal neurons (right) with spiking activity significantly inhibited or excited (*p* < 0.05; Unchanged, *p* >= 0.05; rank sum test) by inhibiting D1 SPN in VLS. Neurons with a mean spike rate above 1 Hz in control trials during the silencing window were analyzed. This includes 25 striatal projection neurons (n = 5 sessions, 5 mice) and 156 ALM neurons (n = 6 sessions, 6 mice). **f.** Schema of the model explaining the data (**f1**). Schema of ALM dynamics during striatal inhibition in the model (**f2**). **g.** ALM activity in a two-dimensional space along the ramp and middle mode during inhibition of D1 SPN in VLS. The mean trajectory of trials with silencing (lick between 1.4 - 1.7 s) is shown. Arrows indicate the direction in which the trajectory evolves. **h.** Left, striatum recording during unilateral D1 SPN inhibition in VLS. Middle and right, similarity matrix of striatal population activity between trial types. Middle, control vs. ref trial (lick between 1.4 - 1.7 s). Right, D1 SPN inhibition trials vs. ref trial. **i.** Points with the peak correlation in the similarity matrix in **h**. Lines, mean. Shades, SEM (hierarchical bootstrap). **j.** Striatal population activity along the ramp mode (n = 10 sessions, 5 mice), trials grouped by lick time. Lines, grand mean. Shades, SEM (hierarchical bootstrap). Left, control. Right, D1 SPN inhibition trials. **k-m.** Same as in **h-j** but for ALM recording during bilateral inhibition of D1 SPN in VLS. n = 6 sessions, 6 mice.

Transient bilateral inhibition of D1 SPN in the ventrolateral striatum (VLS) during the delay epoch (0.6 s duration, starting 0.6 s after the cue onset; 488 nm, 0.25 - 0.5 mW) significantly increased the no-response rate by 38 (5.7 - 41)% (mean, 95% confidence interval, n = 6 mice; Fig. 6cd). In trials where mice licked after the inhibition, the median lick time was shifted later by 1.0 (0.69 - 1.4) s (mean, 95% confidence interval), significantly longer than the duration of photostimulation, and the effect of ALM silencing (Fig. 5).

The shift in lick time caused by unilateral inhibition was approximately half that of bilateral inhibition, suggesting an additive effect of this manipulation (Fig. 6cd, light blue). Inhibiting VLS before the cue did not affect the subsequent lick time distribution, implying that the striatum is specifically involved in lick timing control after the cue and that this manipulation has no long-lasting effect (Fig. 6cd). The behavioral effect of VLS inhibition was stronger than dorsomedial striatum (DMS) inhibition (Extended Data Fig. 8i-l, n = 6 mice), consistent with the strong anatomical and functional connections between VLS and ALM^53–55^.

To measure the impact of D1-SPN inhibition on striatal dynamics, we performed optrode recordings (103 neurons, 5 mice). During unilateral inhibition, population activity patterns comprising all striatal cell types in VLS, appeared to effectively ‘rewind’ their progression: they stopped unfolding, with a slight recession in the points with peak correlation (quantified by similarity matrix, Fig. 6hi). After the inhibition, activity patterns developed from the post-inhibition state, parallel to those in the unperturbed condition (Fig. 6i). During D1-SPN inhibition, striatal population activity patterns stopped unfolding but remained highly correlated with those in the reference unperturbed trials (Fig. 6h). This indicates that D1-SPN inhibition did not deviate striatal population activity patterns from their natural patterns, implying that it exerted an ‘on-manifold’ perturbation of striatal dynamics^43^.

To further quantify the effect of inhibition on striatal dynamics representing time, we analyzed the striatal RM activity and decoded T_to lick_. Unlike ALM silencing, which rapidly decreased the striatal activity at the stimulation onset (Fig. 5j), both striatal RM activity and decoded T_to lick_ gradually decayed during photoinhibition (Fig. 6j and Extended Data Fig. 9l). Because significantly photoinhibited cells (i.e., putative D1 SPN expressing stGtACR1) showed rapid silencing at light onset (within 50 ms; Extended Data Fig. 10l), the gradual decay in timing dynamics is unlikely due to slow photoinhibition but more likely driven by network effects (for example, D1-SPN may modulate thalamic activity via the substantia nigra reticulata, and these thalamic neurons project back to the striatum). After inhibition, the ramp restarted in parallel to control conditions from the post-inhibition level (Fig. 6j and Extended Data Fig. 9i and l). Hence, striatal timing dynamics slowly rewind during unilateral VLS D1-SPN inhibition, as if the D1 SPN inhibition is integrated into the timing dynamics. Altogether, D1 SPN inhibition exerts an on-manifold influence on striatal dynamics (Fig. 1c4), implying that D1 SPN has a highly specific role in implementing the integrator: it may be a part of the integrator or providing on-manifold input to the integrator.

### Striatum is required for ALM timing dynamics

In our network model (Fig. 6f and Extended Data Fig. 1f and 11g), ALM timing dynamics follow those generated by the subcortical integrator. If so, VLS D1-SPN inhibition should rewind ALM timing dynamics as well. To test this hypothesis, we recorded ALM activity during bilateral VLS D1-SPN inhibition (255 neurons, 6 mice). VLS D1-SPN inhibition had relatively minor effects on ALM spiking activity both during ITI and delay epoch: 16.7% of ALM neurons were significantly inhibited and 18.0% were excited during D1-SPN inhibition (Fig. 6e, right). The mean spiking activity decreased by 0.17 ± 0.14 spikes per second (mean ± SEM) during D1-SPN inhibition in the ITI. Thus, VLS D1-SPN is not the major excitatory drive of ALM activity.

Notably, however, VLS D1-SPN inhibition during the delay significantly impacted the timing dynamics in the ALM. During VLS D1-SPN inhibition, ALM population activity patterns paused their progression with a slight recession in the points showing peak correlation (Fig. 6kl). Similar to striatal dynamics, ALM activity patterns did not deviate from natural activity patterns during D1-SPN inhibition, suggesting that D1-SPN inhibition exerted an on-manifold perturbation of ALM dynamics.

ALM RM activity and decoded T_to lick_ gradually decayed during VLS D1-SPN inhibition but resumed ramping in parallel to control trials after inhibition ended, without a rapid recovery phase (Fig. 6m and Extended Data Fig. 9m and p). Consistent with ‘rewinding’, during VLS D1-SPN inhibition, ALM dynamics in the two-dimensional space defined by RM and MM evolved in nearly the opposite direction from the normal trajectory (Fig. 6g). This contrasts with ALM dynamics during ALM silencing, where activity moved toward zero point (See Extended Data Fig. 12 for quantification of rewinding vs. activity moving toward zero). Unilateral VLS inhibition induced a similar, albeit weaker, decay in RM and decoded T_to lick_ in the ipsilateral ALM (Extended Data Fig. 13d-f), similar to how unilateral VLS inhibition affected striatal timing dynamics (Fig. 6j). Thus, while D1-SPN is not a major excitatory drive of ALM activity, it strongly influences timing dynamics in ALM. These results are consistent with a model where ALM timing dynamics follow those generated in the subcortical area (Extended Data Fig. 1f and 11g), rather than ALM maintaining an additional integrator (Extended Data Fig. 1c and 11e).

The impact of ALM and VLS inhibition on ALM timing dynamics differs qualitatively. Even a weak ALM inhibition (0.3 mW instead of the 1.5 mW used in Fig. 5) caused a weak yet rapid decay in RM activity at the onset of photostimulation, followed by a recovery of ramping during photostimulation (Extended Data Fig. 13a-c) and a mild behavioral effect (shifted the median lick time by 0.13 (0.028 - 0.26) s; mean, 95% confidence interval; n = 5 mice). Thus, the gradual decay in ALM timing dynamics during D1-SPN inhibition cannot be explained by its weak inhibitory effect.

Overall, VLS D1-SPN inhibition has a stronger impact on behavior than ALM silencing, despite its weaker effect on mean spike rates. This strong behavioral effect is likely due to its on-manifold impact on timing dynamics. The absence of rapid recovery of timing dynamics after striatal inhibition (Fig. 6m and Extended Data Fig. 9m) suggests there is no other independent area functioning as a timer to restore the activity. These findings support a model where the striatum (and/or subcortical areas situated between the striatum and ALM) implements an integrator that generates timing dynamics, with ALM timing dynamics reflecting those generated by this subcortical integrator (Fig. 6f).

## Discussion

The frontal cortex and striatum often exhibit similar activity patterns and are essential for motor timing and many other behaviors^4–8,19,30^ (Fig. 3). This poses a challenge in distinguishing their functional roles. To address this, we conducted a series of transient perturbations coupled with multi-regional electrophysiology. Across conditions, both ALM and striatum population activity predicted subsequent lick timing even after perturbations (Extended Data Fig. 9), indicating a tight causal link between the dynamics in these brain areas and motor timing. All transient manipulations temporally shifted subsequent timing dynamics in parallel to unperturbed conditions beyond the perturbation period and affected lick time. Furthermore, the extent of the shift depended on the strength of manipulations (unilateral vs. bilateral striatal inhibition, and ALM silencing with different laser powers; Extended Data Fig. 13) as if the perturbation was integrated into the timing dynamics. These findings support the hypothesis that an integrator mediates the generation of timing dynamics.

Importantly, depending on the manipulated brain areas, the multi-regional dynamics during and after perturbation exhibited striking differences: Silencing ALM paused the ‘timer’ without erasing the timing information in the striatum, whereas inhibiting the striatum effectively rewound the ‘timer’ in both areas. Our findings support a model where the subcortical areas (striatum and/or subcortical areas situated between the striatum and ALM) function as an integrator generating timing dynamics in response to inputs provided via ALM (Fig. 5d, 6f, and Extended Data Fig. 1f and 11g). The trial-history information encoded in ALM (Fig. 4) may adjust the ramping slope according to trial history and guide lick timing. ALM follows the ramping activity generated in the subcortical integrator, which is likely critical for ALM to trigger a lick^19^.

The neocortex providing inputs to control the subcortical integrator may be a general mechanism of temporal integration within the cortico-basal ganglia loop. First, singing mice (*Scotinomys teguina*) adjust their song durations based on social context. Silencing the frontal cortex reduces this context-dependent modulation^79^, supporting our model where the frontal cortex adjusts the timer based on contextual information^79,80^. Second, temperature manipulation of the striatum influences the perception of duration^36^, implying that the striatum is essential for both controlling action timing and perceiving time. Third, ramping activity correlated with temporal integration of external signals, or evidence accumulation, is observed in both the frontal cortex and striatum during decision-making tasks, and the striatum is a key site for evidence accumulation^14,30,81–83^. Our network model may explain the mechanism of temporal integration across motor and cognitive behaviors.

Neural correlates and the causality of manipulation on behavior are insufficient to distinguish multi-regional dynamics models (Extended Data Fig. 1 and 11). Specifically, prolonged manipulations that encompass the entire trial (such as muscimol infusion and chemogenetics) have limited ability to differentiate between models (Extended Data Fig. 1, bottom). Therefore, combining transient perturbation with large-scale electrophysiology is critical. This strategy can be employed to dissect multi-regional dynamics across behaviors. Importantly, stringent consideration of behavioral adaptation to perturbations^61,62^, and calibration of photostimulation conditions to prevent rebound are critical for interpretable and reproducible perturbation experiments (Extended Data Fig. 7j-q).

In a memory-guided licking task with a sensory cue signaling proper lick time, transient ALM silencing was followed by a rapid recovery of ramping activity to the original trajectory (akin to Fig. 1c1), regardless of the strength of inhibition^44,45^. In contrast, identical ALM manipulation during the timing task led to a temporal shift in ramping activity, with the extent of the shift depending on the inhibition strength. Thus, depending on the task, the same brain area (ALM) and similar neural activity (ramping activity) are governed by different dynamical systems, presumably to optimize computation for the task at hand.

Similarly, ALM encoded trial history across trials only when mice used trial history to adapt lick timing. We expected that silencing ALM during the ITI would alter the subsequent lick time by affecting this trial history information. Unexpectedly, ALM activity recovered rapidly following ITI silencing and did not affect lick time (Fig. 5a and Extended Data Fig. 9q). This implies that trial history information is robustly maintained across brain areas^63–66^.

In a dynamical system, the evolution of activity states is shaped by both the initial conditions and external inputs of a network^9,31,46,84^. Although we treated trial history information as an input to the integrator in models, it probably functions as both initial conditions and external inputs to determine action timing. Temporal integration can be implemented by tuned feedback loops, and feedforward networks^23,25–27,67–69^. Regardless of the implementation of temporal integration (Extended Data Fig. 1 and 11), our results are consistent with the ALM function as an input/follower to the subcortical integrator. The striatum predominantly comprises inhibitory neurons with relatively sparse lateral connections^70^, so it is unlikely that D1 SPN alone implements an integrator. The striatum indirectly modulates thalamic activity through other basal ganglia nuclei, while intralaminar thalamic nuclei provide direct excitatory input to the striatum^53,54,70,71,85,86^. Future perturbation experiments across areas within this long-range subcortical loop may help elucidate how the integrator is fully implemented in these subcortical areas.

## Acknowledgments

We thank P. Dayan, M. Andermann, N. Li, D. Fitzpatrick, T. Wang, M. Sarvestani, L. Colgan, B. Mensh, R. Heldman, K. Citrin, T. Yiu, X. Yu, and Inagaki lab members for comments on the manuscript, K. Daie, S. Romani and S. Saxena for discussions, P. Scarpinato and N. Spiller for DeepLabCut pipeline, K. Shirley for imaging, and H. Shearin and other MPFI ARC members for animal care. This work was funded by Max Planck Florida Institute for Neuroscience (H.K.I.), Max Planck Free Floater Program (H.K.I.), NIH New Innovator Award (NINDS and OD; 1DP2NS132108; H.K.I.), Searle Scholars Program (H.K.I.), Klingenstein-Simons Fellowship (H.K.I.), and McKnight Scholar Award (H.K.I.). **Author contributions**: Conceptualization: HKI; Investigation: ZY, MI, CG, LF, HKI; Funding acquisition: HKI; Supervision: HKI; Writing: ZY, HKI. **Declaration of interests**: Authors declare that they have no competing interests.

## METHOD DETAILS

### EXPERIMENTAL MODEL AND SUBJECT DETAILS

#### Mice

This study is based on both adult male and female mice (age > P60). We used five mouse lines: C57Bl/6J (JAX# 000664), VGAT-ChR2-EYFP^87^ (JAX #14548), Drd1-cre FK150^74^, Adora2-cre KG126^74^, R26-LNL-GtACR1-Fred-KV2.1^88^ (JAX #33089). See Extended Data Table 1 for mice used in each experiment.

All procedures were in accordance with protocols approved by the MPFI IACUC committee. We followed the published water restriction protocol^89^. Mice were housed in a 12:12 reverse light: dark cycle and behaviorally tested during the dark phase. A typical behavioral session lasts between 1 and 2 hours. Mice obtained all of their water in the behavior apparatus (approximately 0.6 ml per day). Mice were implanted with a titanium headpost for head fixation^89^ and single-housed. For cortical photoinhibition, mice were implanted with a clear skull cap^48^. For bilateral D1/D2 SPN silencing, tapered fiber optics (1.0 mm taper, NA 0.37, core diameter 200 µm, Doric lenses) were bilaterally implanted during the headpost surgery around the following target coordinates (Extended Data Fig. 6c; Bregma, mm): AP −0.3, ML ±3, DV 3.5 for the ventral lateral striatum (VLS); AP 0.6, ML ±1.5, DV 3 for the dorsal medial striatum (DMS). Craniotomies for recording were made after behavioral training.

#### Viral injection

To virally express stGtACR1 in the striatum (Extended Data Fig. 8q), we followed published protocols (dx.doi.org/10.17504/protocols.io.bctxiwpn) for virus injection. AAV2/5 CamKII-stGtACR1-FusionRed (titer: 9.5×10^12) was injected into AP −0.3 mm, ML 3 mm, DV 2.75 and 3.5 mm, 100nl each depth. The same tapered fiber optics described above were bilaterally implanted at DV 3.5 mm.

#### Behavior

At the beginning of each trial, an auditory cue was presented, which consisted of three repeats of pure tones (3 kHz, 150 ms duration with 100 ms inter-tone intervals, 74 dB). A delay epoch started from the onset of the cue presentation. Licking during the delay epoch aborted the trial without water reward, followed by a 1.5 s timeout epoch. Licking during the 10 s answer epoch following the delay was considered a ‘correct lick’, and a water reward (approximately 2 µL/drop) was delivered immediately, followed by a 1.5 s consumption epoch. If mice did not lick during the 10 s answer period, the trial would end without a reward. Trials were separated by an inter-trial interval (ITI) randomly sampled from an exponential distribution with a mean of 3 s with 1 s offset (with a maximum ITI of 7 s). This prevented mice from predicting the trial onset without cue. Animals had to withhold licking during the full ITI epoch for the next trial to begin (otherwise the ITI epoch repeated). In approximately 10% of randomly interleaved trials, the auditory cue was omitted to assess spontaneous lick rate (‘no cue’ trials). No water reward was delivered in no cue trials.

We followed the protocol described in Majumder et al, 2023^90^ for training. In brief, the delay duration increased from 0.1 s to 1.8 s gradually based on the animal’s performance^90^. Once mice reached 1.8 s delay, we started either the switching delay (Extended Data Fig. 2a), the random delay (Extended Data Fig. 2b), or the constant delay conditions (Extended Data Fig. 2c). In the switching delay condition, we switched delay between 1 vs. 3 s or 1 vs. 1.8 s every 30 - 70 trials (the number of trials was randomly selected from 30-70 and not contingent upon behavior). Similarly, in the random delay condition, we randomly switched delay among 0.5, 1.0, 1.5, 2.0, 3.0, or 5.0 s every 30 - 70 trials. For the constant delay condition, mice were trained with a constant delay of 1.5 s across sessions for at least two weeks. Otherwise, the task design and reward contingency remained the same. ALM and striatal perturbation experiments (Fig. 5 and 6) were performed under the switching delay condition. To avoid human bias, the behavior was automatically controlled by Bpod (Sanworks) and custom MATLAB codes.

#### Optogenetics

Photostimulation was deployed on < 25% in randomly selected trials. To prevent mice from distinguishing photostimulation trials from control trials using visual cues, a ‘masking flash’ (1 ms pulses at 10 Hz) was delivered using 470 nm LEDs (Luxeon Star) throughout the trial. For both ChR2 and stGtACR1, we used a 488 nm laser (OBIS 488 - 150C, Coherent).

The ChR2-assisted photoinhibition of the dorsal cortices was performed through clear-skull cap^48^ (Fig. 2e) or craniotomy (in case of simultaneous recording; Fig. 5). We scanned the 488 nm laser light using Galvo mirrors. We stimulated GABAergic interneurons in Vgat-ChR2-EYFP mice starting at 0.6 s after the cue, lasting for 1.2 s (including 0.2 s ramping down; Fig. 2e) or 0.6 s duration (including 0.3 s ramping down; Fig. 5). Time-averaged laser power was 1.5 (or 0.3 for Extended Data Fig 13) mW per spot (8 spots in total: 4 spots in each hemisphere centered around the target coordinates with 1 mm intervals; We photoinhibited each spot sequentially at the rate of 5 ms per step). For Fig. 2e, the targeted brain area was randomly selected for each photostimulation trial. The target coordinates were AP 2.5, ML ±1.5 (ALM); AP 0.5, ML ±1.5 (M1B); AP 0.5, ML ±2.5 (S1TJ); AP −1.0, ML ±1.5 (S1TR); AP −1.0, ML ±3.0 (S1B); AP −2, ML ±1.5 (PPC); and AP −2.5, ML ±3.5 (V1) respectively (Bregma, mm).

To silence D1 or D2 expressing SPNs using stGtACR1 (Fig. 6), we delivered photostimuli (0.25 or 0.5mW, 488 nm) bilaterally (Fig. 6k-m) or unilaterally (in case of optrode; Fig. 6h-j) in the striatum starting 0.6 s after the cue and lasting for 0.6 s (including 0.3 s ramping down). The light was delivered through implanted fiber optics and intensity was measured at the fiber tip.

#### Extracellular electrophysiology

A small craniotomy (diameter, 0.5 - 1 mm) was made over the recording sites one day before the first recording session. Extracellular spikes were recorded acutely using 64-channel two-shank silicon probes (H-2, Cambridge Neurotech) for ALM, and Neuropixels probe 1.0^56^ for striatum. For the H-2 probes, voltage signals were multiplexed, recorded on a PCI6133 board (National instrument), and digitized at 400 kHz (14-bit). All recordings were made with the open-source software SpikeGLX (http://billkarsh.github.io/SpikeGLX/). During recordings, the craniotomy was immersed in a cortex buffer (125 mM NaCl, 5 mM KCl, 10 mM glucose, 10 mM HEPES, 2 mM MgSO_4_, 2 mM CaCl_2_; adjust pH to 7.4). Brain tissue was allowed to settle for at least five minutes before recordings.

For the optrode recordings (Fig. 6h-j), we used 64-channel two-shank silicon optrodes with a 1.0 mm taper fiber optic attached adjacently (NA 0.22, core diameter 200 µm, Cambridge Neurotech). Optrode was acutely inserted each session and the light delivery protocol was identical to that used for behavioral experiments described in ***Optogenetics***. Neuropixels probe and optrode tracks labeled with CM-DiI were used to determine recording locations^91^.

#### Histology

Mice were perfused transcardially with PBS, followed by 4% PFA / 0.1 M PBS. To reconstruct recording tracks, we either generated coronal sections followed by conventional imaging (protocol described in *Inagaki et al, 2022*^92^), or cleared the brain followed by light-sheet microscopy. To clear the brain, we used the EZ Clear method^93^. We followed the previous protocol to map the recording tracks to Allen Common Coordinate Framework (CCF)^91^. Extended Data Fig. 6a is based on a brain imaged in Guo et al, 2017^94^.

### QUANTIFICATION AND STATISTICAL ANALYSIS

#### Behavioral analysis

We analyzed the time of the first lick after the cue onset in each trial. Lick time was measured by detecting the tongue’s contact with the lick port using an electrical lick detector. For optogenetic experiments (Fig. 5ab, 6cd, and Extended Data Fig. 8), we analyzed trials with the first lick occurring after the onset time of photostimulation (0.6 s after the cue) in both control and photostimulated trials to compare the effect of photostimulation on behavior. The no-response rate (Fig. 6d, right, and Extended Data Fig. 8) was calculated as the probability of mice not responding within 5 s after the cue. The shift in lick time (Δ lick time; Fig. 5b, 6d, and Extended Data Fig. 8) was based on the median lick time. Post-stim lick rate (Extended Data Fig. 7bc) was calculated as the probability of mice licking within 0.6 s after the photostimulation offset time in no cue trials. To analyze behavior while the mice were engaged in the task, we analyzed all trials between the first occurrence of five consecutive cue trials with licks and 20 trials before the last occurrence of three consecutive no-response trials.

To analyze whether optogenetic manipulation affects the separation of lick time distributions between different delay blocks in the switching delay condition (short vs. long delay blocks; Extended Data Fig. 8d, h, l, and p), we performed a receiver operating characteristic (ROC) analysis. First, we conducted ROC analysis to distinguish lick time distributions between the two delay blocks (for control and photostimulation trials separately). We then quantified the area under the curve (AUC) to measure the separation in lick time distributions between delay blcoks, and compared these values between control and photostimulation trials.

Due to the attenuation of behavioral effects of optogenetic manipulation (Extended Data Fig. 7f-i), we restricted analyses of both behavioral and physiological data to the first (for striatal manipulation) or the first two (for ALM manipulation) manipulation sessions per mouse. All analyses, including the calculation of confidence intervals and p-values, were performed using a hierarchical bootstrap, unless stated otherwise. First, we randomly selected animals with replacements. Second, we randomly selected sessions for each animal with replacement. Third, we randomly selected trials for each session with replacements. Then, we calculated the behavioral metrics described above. This procedure was repeated 1000 times to estimate the mean, confidence intervals, and statistics.

##### Behavioral simulation

We estimated the optimal lick time in the timing task (Extended Data Fig. 2g-k). Based on the mean and standard deviation (std) of the lick time distribution in the data (std lick time = −0.2 + 1.1 × mean lick time; Extended Data Fig. 2f), and assuming an inverse Gaussian distribution of lick time^95–97^, we randomly sampled hypothetical lick times in 1000 trials for a given mean lick time. Then, following the task structure (identical to that described in ***Behavior***), we determined whether the agent would receive water or not for each trial and calculated the estimated reward amount per trial or time.

##### Trial-history regression analysis

For the linear regression analysis in Fig. 2d and Extended Data Fig. 2l, we tested 42 combinations of regressors with 1 - 6 lags with 5-fold cross-validation. The median absolute deviation (MAD) of lick time explained by different regression models was calculated as 1 - R1 / R2, where R1 is the median of the absolute value of the model residuals, and R2 is the median of the absolute value of the null model residuals.

#### Videography analysis

High-speed (300 Hz) videography of orofacial movement (side view) was acquired using a CMOS camera (Chameleon3 CM3-U3-13Y3M-CS, FLIR) with IR illumination (940nm LED). We used DeepLabCut^98^ to track the movement of the tongue and jaw. Movements along the dorsoventral direction were analyzed and plotted in Extended Data Fig. 3. Trajectories were normalized by subtracting the mean position before the cue from each trajectory and then dividing by the minimum value within a session (thus, the downward movement of the tongue and jaw looks upward in the plot). The onset of jaw movement in each trial is the first time point after the cue when the normalized movement trajectory exceeds 10% of the max value. The onset of tongue movement is when DeepLabCut first detects the tongue after the cue. In two out of 34 mice, mice moved jaws within 200 ms after the cue in some trials. These trials were excluded from the analysis for the average jaw and tongue onset analyses (Extended Data Fig. 3d), as these rare early subthreshold movements are likely startled responses to the cue.

#### Extracellular recording analysis

##### Spike sorting and cell type classification

JRClust^99^ (https://github.com/JaneliaSciComp/JRCLUST) with manual curations was used for spike sorting. We used quality metrics (described in Majumder et al, 2023^90^) to select single units. Units with a total trial number of less than 75 were excluded from analyses.

For ALM recording, units with a mean spike rate above 0.5 Hz were analyzed. This includes 4467 putative pyramidal neurons (spike width >= 0.5 ms^48^) out of a total of 5093 neurons. For striatal recording, units within the striatum (regions annotated as “striatum”, “caudoputamen”, and “fundus of striatum” after registration to the Allen CCF) with a mean spike rate above 0.1 Hz were analyzed. This contains 1217 striatal projection neurons (spike width >= 0.4 ms and with post-spike suppression duration <= 40 ms), 584 fast-spiking interneurons (spike width < 0.4 ms and with less than 10% chance of having a long interspike interval), 127 tonically active neurons (TAN), and 44 other unidentified interneurons^57^. For the single session analyses (decoding and projection to modes), only putative pyramidal neurons were analyzed for the ALM recording, while all neurons were included for the striatal recording data. See Extended Data Table 1 for the number of recorded neurons in each experiment.

##### Similarity matrices

To plot the similarity matrix of population activity, we calculated the mean spike activity of individual neurons across trials with different lick time ranges to yield a population activity matrix, with the number of rows equal to the number of neurons and the number of columns equal to the number of time points (200 ms bin). For Fig. 3, we calculated pairwise Pearson’s correlation of these population activity matrices between trials with lick times between 1.40 - 1.55 s (reference trials) and the trials with other lick time ranges. For Fig. 5 and 6, we compared the pairwise Pearson’s correlation between unperturbed trials with lick times between 1.4 - 1.7 s (reference trials) and the photostimulation trials with lick times between 1.7∼ 2.0s). As a control, we subselected unperturbed trials with lick times closest to the median lick time in the unperturbed condition (the number of trials was matched to the number of trials as in the photostimulation condition). The choice of reference trials did not change results qualitatively. For each similarity matrix, we identified the points along the Y-axis with the maximum correlation (above 0.8) for each time point, and repeated this procedure with the hierarchical bootstrap (Fig. 3d, 3i, 5f, 5i, 6i and 6l).

##### Single-cell analyses

To plot the peri-stimulus time histogram (PSTH) of example cells, PSTHs were calculated based on 1 ms time bin, and smoothed with a 200 ms causal boxcar filter unless specified otherwise. To temporally warp PSTH for individual cells, we linearly scaled the spike timing after the cue, based on the time from cue to lick. Specifically, Spike time_warped_ = Spike time_original_ / (LT_trial to be warped_ / LT_target warp time_), where LT denotes the first lick time in each trial, and LTtarget _warp time_ = 1 s.

Across-trial variance (Extended Data Fig. 4a-c) was calculated as the variance of spiking activity across trials for the original or temporally warped data (the across-trial variance was calculated for five 200 ms time windows after the cue and then averaged).

To quantify the number of cells that significantly increase/decrease spike rate before the lick compared to baseline, trial-averaged spike rate of 0.2 - 0.5 s before the lick was compared with those 0 - 1 s before the cue. Signed rank tests were performed to determine if the spike rate difference was significant.

To calculate the proportion of cells affected/unaffected by photostimulation (Fig. 5c, 6e, and Extended Data Fig. 10), we analyzed the spikes within the time window of 50 - 250 ms from the photostimulation onset time. To quantify the effect of photostimulation, trials with licks before the photostimulation onset time were excluded from the analysis. For individual cells, the spike rate in control and photostimulation trials was compared using the rank sum test. Cells with a mean spike rate above 1 Hz during this window and more than 10 trials per condition were analyzed.

In Extended Data Fig. 4, we analyzed the partial rank correlation between the spike rate (in specific time windows) and the lick time in previous trials, removing the effect of upcoming lick time, for each cell (we only analyzed trials after rewarded trials to avoid confound caused by the representation of rewards; analysis of previous unrewarded trials yielded similar results). Specifically, we calculated the rank correlation between spike rate (R) vs. previous lick time (P) (*ρRP*), rank correlation between R vs. upcoming lick time (U) (*ρRU*), and rank correlation between P and U (*ρPU*). Then the partial correlation between spike rate vs. lick time in the previous trial removing the effect of upcoming lick time is as follows:

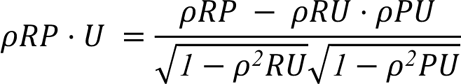

As controls, we performed a trial shuffle test, which shuffles the trial order and destroys trial history, and a session permutation test to avoid the confound of nonsensical correlations (1000 iterations)^58^. The proportion of cells with a correlation higher than the chance level estimated by these controls is shown.

##### Dimensionality reduction

We characterized population activity patterns between the cue and the lick by defining modes that differentiate the baseline activity during the inter-trial interval (ITI; 0 - 1 s before the cue) from the activity during specific 300 ms time windows after the cue: 0 - 0.3 s after the cue (cue mode, **CM**), 0.5 - 0.8 s before the lick (middle mode, **MM**), 0.2 - 0.5 s before the lick (ramp mode, **RM**), and 0 - 0.3 s after the lick (execution mode, **EM**).

Specifically, to calculate **RM** for a population of *n* recorded neurons, we looked for an *n* × 1 unit vector that maximally distinguished the mean activity before the trial onset (0 - 1 s before cue; ***r***_before cue_) and the mean activity before the first lick (0.2 - 0.5 s before the first lick; ***r***_before lick_) in the *n-dimensional* activity space. We defined a population ramping vector: ***w*** = ***r***_before lick_ – ***r***_before cue_. **RM** is ***w*** normalized by its norm. Similarly, we defined **CM, MM**, **EM** using different time windows, and **MM** was orthogonalized to **RM**, and **CM** was orthogonalized to both **MM** and **RM** using the Gram-Schmidt process. Thus, the upper limit of the sum of variance explained (**CM**+**MM**+**RM**) in Extended Data Fig. 5 is 1. **EM** was orthogonalized to **RM** (Extended Data Fig. 4j3 and k3).

To calculate the trial-history mode, we first calculated the rank correlation between the ITI activity (0 - 1 s before the cue) and the predicted lick time across trials for each neuron. To predict the lick time for each trial, we used the linear regression model described in Fig. 2d following 5-fold cross-validation. We obtained an n × 1 unit vector representing the rank correlation of each neuron and normalized it by its norm to calculate the trial-history mode.

In Fig. 3, 4, and Extended Data Fig. 4 and 5, we have pooled cells recorded across sessions (i.e., pseudo-sessions). For each cell, we randomly selected 50 unperturbed control trials to define the mode. These unperturbed trials met the following criteria: the first lick occurred within 1 to 3 s after the cue, and there were no licks 3 s before the cue onset. Then, we selected a different set of trials to project the activity along these modes. Only neurons with more than 10 trials within all six lick time ranges were included. The six lick time ranges are: 0.80 - 1.10, 1.10 - 1.25, 1.25 - 1.40, 1.40 - 1.55, 1.55 - 1.70, and 1.70 - 2.00 s.

To calculate the variance of spiking activity explained (variance explained) by individual modes (Extended Data Fig. 5), we calculated the squared sum of the activity along individual modes after subtracting the baseline activity (0 - 0.2 s before the cue), and then divided that by the squared sum of the spike rate across neurons after subtracting the baseline activity. To calculate the variance explained by the sum of **CM**, **MM**, and **RM** reported in the main text, we calculated the variance explained between 0.2s from cue (around when task modulation started) to lick for individual lick time ranges and then averaged across them. We calculated the variance explained by trial-history mode activity similarly but without subtraction of the baseline activity (Extended Data Fig. 4j-l). In Extended Data Fig. 4m, we performed a linear regression analysis between trial-history mode activity during 0 - 1 s before the cue and the upcoming lick time for each iteration of the hierarchical bootstrap. We then plotted the distribution of the linear regression coefficient (slope) across these iterations as a cumulative distribution function.

##### Single session analyses

For single-session analyses (Fig. 5g, and j, 6g, j, and m, Extended Data Fig. 9, 12, and 13), sessions with more than 300 trials and five neurons were analyzed. Spiking activity was binned per 50 ms time window. Activity between 1 s prior to the cue and the first lick in each trial was analyzed (i.e., post-lick activity was excluded as we focused on timing dynamics prior to the first lick). Dimensionality reduction was performed in the same manner as in the pseudo-session analysis, but modes were defined individually for each session. For plots, lick time ranges that exist in at least two-thirds of the analyzed sessions were shown. Therefore, the plotted lick time ranges vary depending on the manipulation conditions.

To decode the time to lick (T_to lick_) from simultaneously recorded neural population activity, we conducted a k-nearest neighbor (kNN) regression analysis. Within each experimental session, trials were partitioned into two sets: a test set comprising randomly selected 100 unperturbed trials and all perturbed trials, and a training set consisting of the remaining trials. For each moment in a test trial (50 ms window), we searched all time points in the training set to identify k data points with the most similar population activity patterns (Mahalanobis distance based on the top principal components explaining 90% of variance). To estimate the time to lick of the test set, we averaged the time to lick in these k nearest neighbors. We tested “k” values between 20 - 50 (which are close to the square root of the number of data points in the training dataset) and found they yielded similar results and did not change conclusions (data not shown). In the paper, we report the results with k = 30. Some sessions showed low decodability due to a small number of recorded neurons, trials, and/or lack of task-modulated cells (Extended Data Fig. 6g). We analyzed sessions in which the kNN decodability (Pearson’s correlation between decoded lick time at the perturbation onset time, i.e., 0.6 s after the cue vs. actual lick time) was higher than 0.25.

To analyze the effect of perturbations systematically, we compared unperturbed vs. perturbed trials after matching the number of trials and decoded time at the perturbation onset time (Extended Data Fig. 9a, c-d, e, g-h, i, k-l, m and o-q, and 13c and f). Specifically, we randomly resampled animals, sessions, and trials hierarchically (hierarchical bootstrap; 1000 iterations). For each perturbed trial in each bootstrap iteration, we identified an unperturbed trial within the same session with the closest decoded time at the perturbation onset time (0.6 s after the cue). Then, we pooled these trials. This procedure allowed us to examine how decoded time (and projection along each mode) changed after the perturbation in conditions where their activity patterns were similar before the perturbation.

For the two-dimensional plots and vector field analysis (Extended Data Fig. 12), we analyzed how activity evolves in the two-dimensional space defined by RM and MM. Spiking activity was binned in 50 ms time windows, and activity between the cue and the first lick in each trial was analyzed. For each session, we projected the activity of ALM neurons along RM and MM. The projection was normalized by the standard deviation of activity among control trials but was not subtracted by the mean so that 0 represents 0 spike activity. For individual activity state (x) in control trials, we calculated the vector 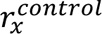 representing the direction activity evolves in the next time point (50 ms time bin) in the two-dimensional state. Then, we calculated the mean vector for individual states in the two-dimensional space by averaging all vectors within a spatial bin of 0.5 along both the MM- and RM-axes (if the spatial bin contains more than 30 data points): 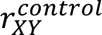, where X and Y denotes the location of the state along MM and RM axes, respectively Similarly, we acquired the vector field during inhibition by pooling all time points during inhibition (100 ms - 400 ms from the inhibition onset) in photostimulation trials to acquire 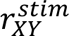. Then, we calculate the direction between 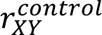 and 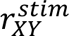 for all states where both control and stim vectors exist. We have excluded points where 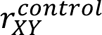 is within *π*/*6* from tanh(Y/X) because if the activity is evolving against the zero point under control conditions, we cannot distinguish between whether during inhibition the activity is rewinding or moving toward the zero point.

#### Network models

Using a dynamical systems approach, we consider four variables representing the average membrane currents (*h*) and spike rates (*r = f(h)*, where *f(h)* is the neural activation function) of neuronal populations in ALM and striatum. Conceptually, in these models, the striatum represents both connections within the striatum and the subcortical loop via the thalamus, which is why there are excitatory connections. In these models, the membrane potential of neuron i, *ℎ*_*i*_(t), was governed by the following nonlinear differential equation:

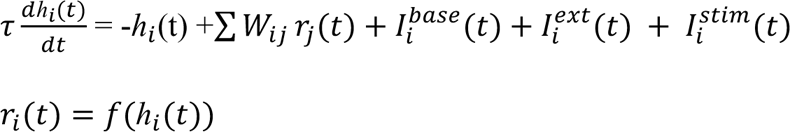

Where τ is the membrane time constant (10 ms), *Wij* is the element of the connectivity matrix between presynaptic neuron j and postsynaptic neuron i, 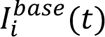 is the baseline input current, 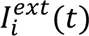 is the external input current, and 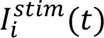 is the negative current mediated by optogenetics to neuron i. The membrane current *ℎ*_*i*_(t) was converted to the spike rate by applying a threshold-linear activation function *f(h) = max(h,0)*.

For integrators mediated by a positive feedback loop (Extended Data Fig. 1), we modeled two neurons in each brain area. The baseline input currents were chosen so that the system displays a stable fixed point at a low spike rate (lower attractor) with a spike rate of 5 spikes per second, consistent with the baseline firing rate observed in the experimental data. The connectivity matrix *W* and the external input *I*^*ext*^(*t*) are shown in Extended Data Fig. 1. In these models, temporal integration is mediated by a continuous attractor, achieved by having an eigenvalue of 1 in the connectivity matrix. For each area, we defined the ramp mode using the same criteria as in the experimental data, and we then plotted spike rate activity along the ramp mode (Extended Data Fig. 1).

We tested models with different connectivity matrices reflecting distinct computational roles of ALM and striatum (Extended Data Fig. 1). In the *externally driven* model (Extended Data Fig. 1a), ALM received a ramping input that scaled with the desired lick times, progressively shifting the location of the fixed point in time. In the *distributed* model (Extended Data Fig. 1b), integration was achieved only when interareal connections between ALM and striatum exists; in the absence of these long-range connections, neither ALM nor striatum displayed slow temporal dynamics. Conversely, in the *redundant* model, ALM and striatum implemented two identical integrators (Extended Data Fig. 1c). Although weakly connected, their behavior was independent of each other’s input. In the *specialized ALM integrator* model (Extended Data Fig. 1d), ALM served as the integrator while the striatum followed ALM dynamics. In the *specialized ALM leaky integrator* model (Extended Data Fig. 1e), ALM integrated the input with substantial leakiness. While this model replicated the rewinding effect of striatal inhibition, it failed to reproduce the effect of ALM silencing. The model that best matched the neural dynamics observed in our data featured the striatum as a perfect integrator and the ALM as a crucial input region (*specialized striatum integrator*, Extended Data Fig. 1f).

To mimic transient perturbation experiments, the negative current was introduced at 0.6 s after the cue and lasted for 0.6 s, including a 300 ms ramp down. For prolonged perturbations, the negative current was applied throughout the trial without a ramp-down. To simulate ALM silencing, both ALM neurons (A1 and A2 in Extended Data Fig. 1) received a negative current *I*^*stim*^(*t*) = −10. To stimulate D1-SPN inhibition, we injected a negative current into one of the striatum neurons (S1). To maintain similar perturbation effects on striatal ramp mode activity across different conditions, the negative current for D1-SPN inhibition was varied across models as follows: −0.3, −0,1. −0.1, −0.2, −0.02, and −0.03 for Extended Data Fig. 1a-f, respectively.

For integrators mediated by recurrent networks with feedforward connections (Extended Data Fig. 11), we modeled four neurons in each area. In these models, recurrent connections in each recurrent stage are not strong enough to generate ramping activity from a step input, and FF connections between stages are essential to amplify the slow time constant and generate ramping activity. To implement temporal scaling, we provided a global inhibition to neurons in the FF network, i.e., 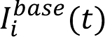 was set to negative. This allows activity to propagate from one neuron to another when the effect of excitatory input exceeds this inhibition. Consequently, the speed of dynamics is controlled by the strength of step input into the network. The connectivity matrix *W* is shown in Extended Data Fig. 11. Transient perturbations were simulated similarly to the positive feedback network. For ALM complete silencing *I*^*stim*^(*t*) = −10 was injected into all ALM neurons. To simulate D1-SPN inhibition, a negative current was injected into half of the striatal neurons (s2 and s3; the result did not change regardless of the choice of two inhibited neurons). To maintain similar perturbation effects on striatal ramp mode activity across different conditions, the negative current for D1-SPN inhibition was varied across models as follows: −5, −5, −100, and −2.5 for Extended Data Fig. 11d-g, respectively. For each area, we defined the ramp and other modes using the same criteria as in the experimental data, and we then plotted spike rate activity along these modes.

#### Statistics

The sample sizes are similar to the sample sizes used in the field. No statistical methods were used to determine the sample size. During spike sorting, experimenters could not tell the trial type and, therefore, were blind to conditions. All *signed rank* and *rank sum* tests were two-sided. All bootstrap was done over 1,000 iterations.

#### Reagent and data availability

The recording data in NWB format will be shared on DANDI at the time of publication. Codes will be available at https://github.com/inagaki-lab/Yang_et_al_2024.

---

**Extended Data Fig 1.**
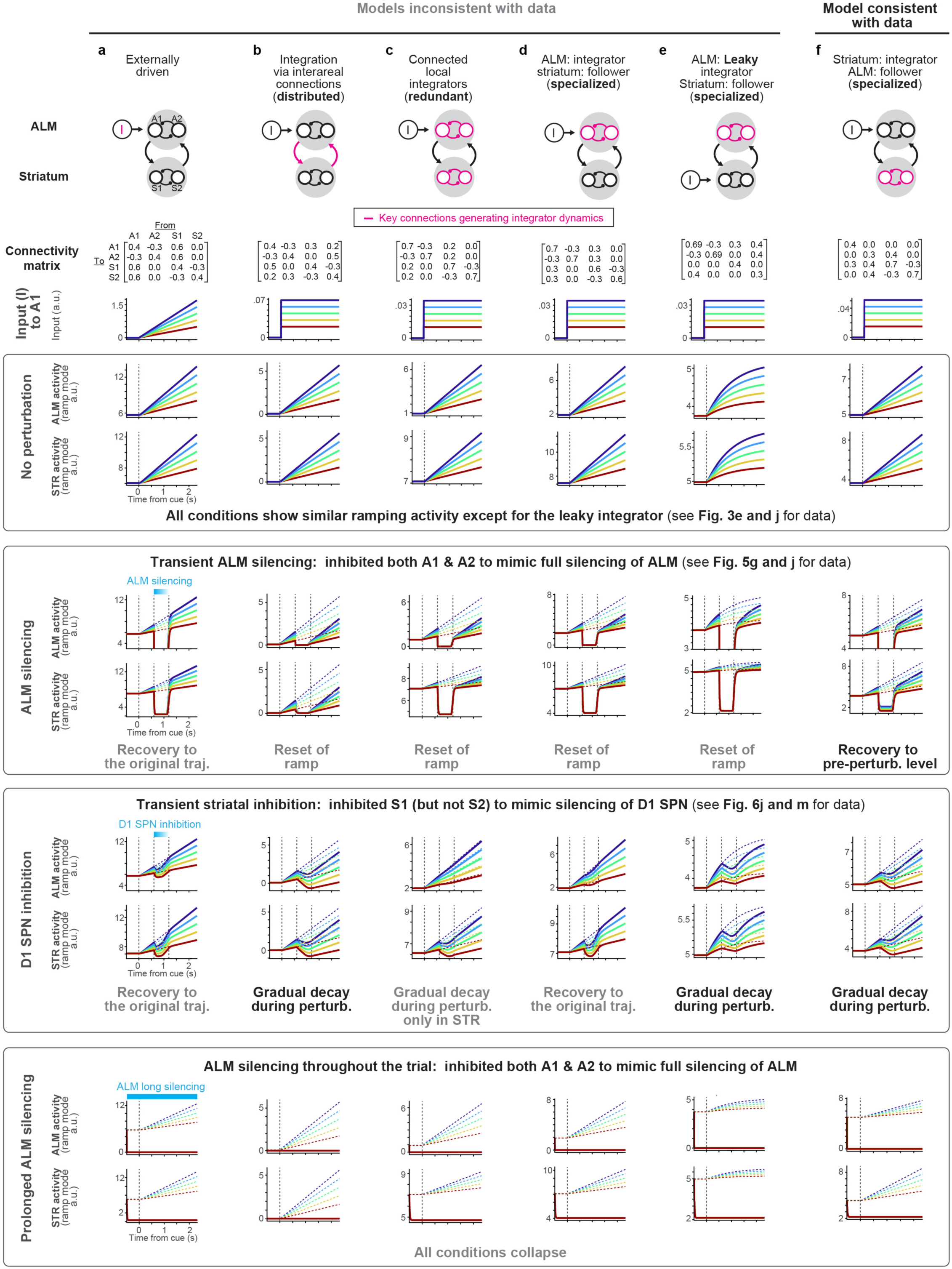
Two-regional network models replicating the ramping activity in ALM and striatum. Multi-regional network models of ALM and striatum: both regions contain two neurons and exhibit ramping activity with temporal scaling as in Fig. 3. Additionally, step input mimicking the trial-history mode activity is provided to ALM (except for **e**; most models yield consistent results even if we provide the input to the striatum, and thus only one configuration is shown). The network configuration, i.e., the connectivity matrix, varies across models, leading to a different location(s) of the integrator(s) and various responses to transient perturbations. ALM silencing was implemented by silencing both neurons in ALM, and striatal inhibition was implemented by silencing one of the striatal neurons (mimicking the silencing of D1 SPNs). Note that prolonged silencing does not distinguish between models (bottom row), underscoring that transient perturbation with concurrent multi-regional recording is essential to differentiate models. **a.** Externally driven model. 1st row, schema of the model, and the connectivity matrix used for the simulation. 2nd row, input (I) into the network. 3rd and 4th row, ALM and striatal ramp mode activity without perturbation. 5th and 6th row, ALM and striatal ramp mode activity during transient ALM silencing (the ALM activity along the ramp mode was cropped for visualization purposes. In all cases, ALM activity during ALM silencing decreased to 0). 7th and 8th row, **ALM** and striatal ramp mode activity during transient striatum inhibition. 9th and 10th row, **ALM** and striatal ramp mode activity during prolonged ALM silencing throughout the trial. Dashed lines, control conditions overlaid. **b.** Same as in **a** but for integration via interareal connections (distributed) model. **c.** Same as in **a** but for coupled local integrators (redundant) model. **d.** Same as in **a** but for ALM being the integrator and striatum being the follower (specialized) model. **e.** Same as in **a** but for ALM being a leaky integrator and striatum being the follower (specialized) model. This model replicates the rewinding of ALM and striatum dynamics during striatal inhibition (where rewinding is caused by the loss of input to the leaky integrator) but cannot reproduce the effect of ALM silencing. **f.** Same as in **a** but for striatum being the integrator and ALM being the follower (specialized) model that replicates the data.

**Extended Data Fig 2.**
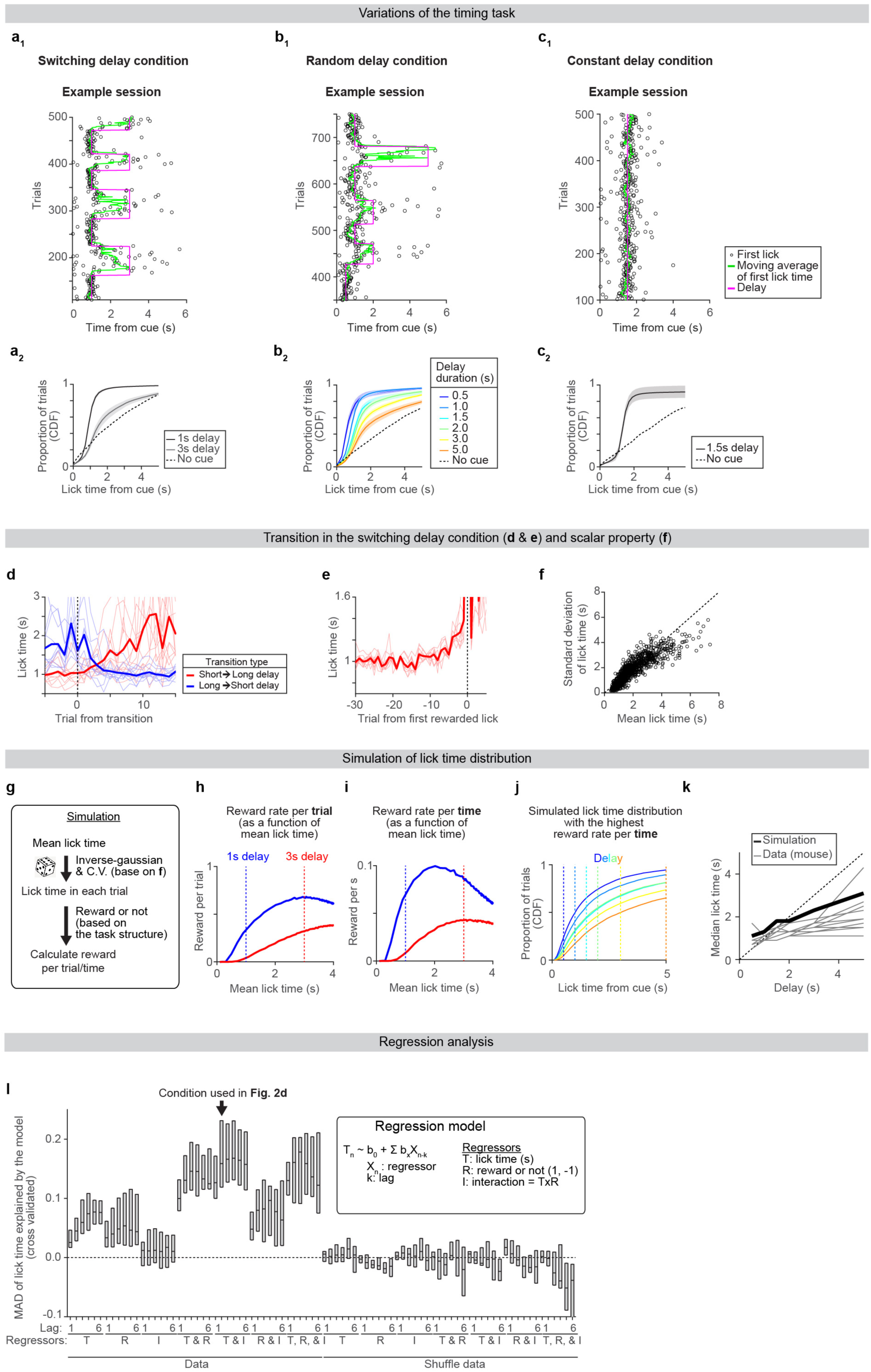
Characterization of the lick time distribution in the lick-timing task. We either changed the delay duration in blocks of trials (**a**, switching delay condition, when switching between two delays; **b**, random delay condition, when switching across multiple delays) or kept the delay identical across sessions/trials (**c**, constant delay condition). When delays switched, mice adjusted their lick times within 10 trials (**d, e**). The lick time distribution exhibited scalar properties, similar to many other timing tasks across species (**f**). Mice often licked earlier than the delay duration, not maximizing reward per trial. However, this lick time distribution (especially in the best-performing mice) is close to the simulated distribution that maximizes the reward amount per time, given the short inter-trial interval and the inverse normal distribution of lick times^95–97,100,101^ (**g-k**, Methods). 42 regression models were screened to identify the best model explaining the lick time (**l**). **a.** Lick time distribution under switching delay condition. Example session (**a1**). Cumulative distribution of lick time in 1 s and 3 s delay blocks (**a2**). Duplicated from Figure 2bc for comparison. **b.** Same as in **a** but for the random delay condition. n = 276 sessions, 17 mice. **c.** Same as in **a** but for the constant delay condition. n = 71 sessions, 13 mice. **d.** Change in lick time after transitioning between 1 and 3 s delay blocks. 0, last trial before delay transition. Thick lines, the mean across mice. Thin lines, individual mice (n =10 mice). **e.** Change in lick time before the first rewarded lick following transitions from a short to a long delay. Thick line, the mean across mice. Thin lines, individual mice (n = 10 mice). **f.** Relationship between the mean and the standard deviation of lick time. Circles, individual sessions (n = 153 sessions, 30 mice). The mean and standard deviation of lick time are correlated, consistent with scalar properties reported across species^102^. **g.** Simulated lick time distribution following an inverse-Gaussian distribution and coefficient of variation (CV) in **f**. Subsequently, based on the task structure, we calculated the reward rate (Methods). **h.** Simulated reward rate per **trial** as a function of mean lick time in 1 s delay block (blue) and 3 s delay block (red). **i.** Simulated reward rate per **time** as a function of mean lick time. Note that the peak reward rate is attained with a shorter mean lick time compared to that in **h**. **j.** Following the procedure described in **g** and **i**, we estimated the optimal mean lick time that yielded the highest reward rate per unit of time for each delay duration (delay duration is indicated by colored vertical dotted lines). We plotted the distribution of simulated lick times in these conditions. Note that in a large proportion of trials, the licks occurred before the end of the delay period, replicating what was observed in the data. **k.** Thick line, the median lick time of simulated optimal lick time distribution in **j**. Thin lines, experimental data (individual mice). The optimal lick time appears to align with the upper limit of the experimental data. **l.** Median absolute deviation (MAD) of lick time explained by different trial-history regression models following cross-validation under the random delay condition (n = 276 sessions, 17 mice; results were consistent in the switching delay condition). Regressors and lags in each model are indicated at the bottom. Arrow, the condition that best explained the data (Fig. 2d). The central line in the box plot, median. Top and bottom edges, 75% and 25% points.

**Extended Data Fig 3.**
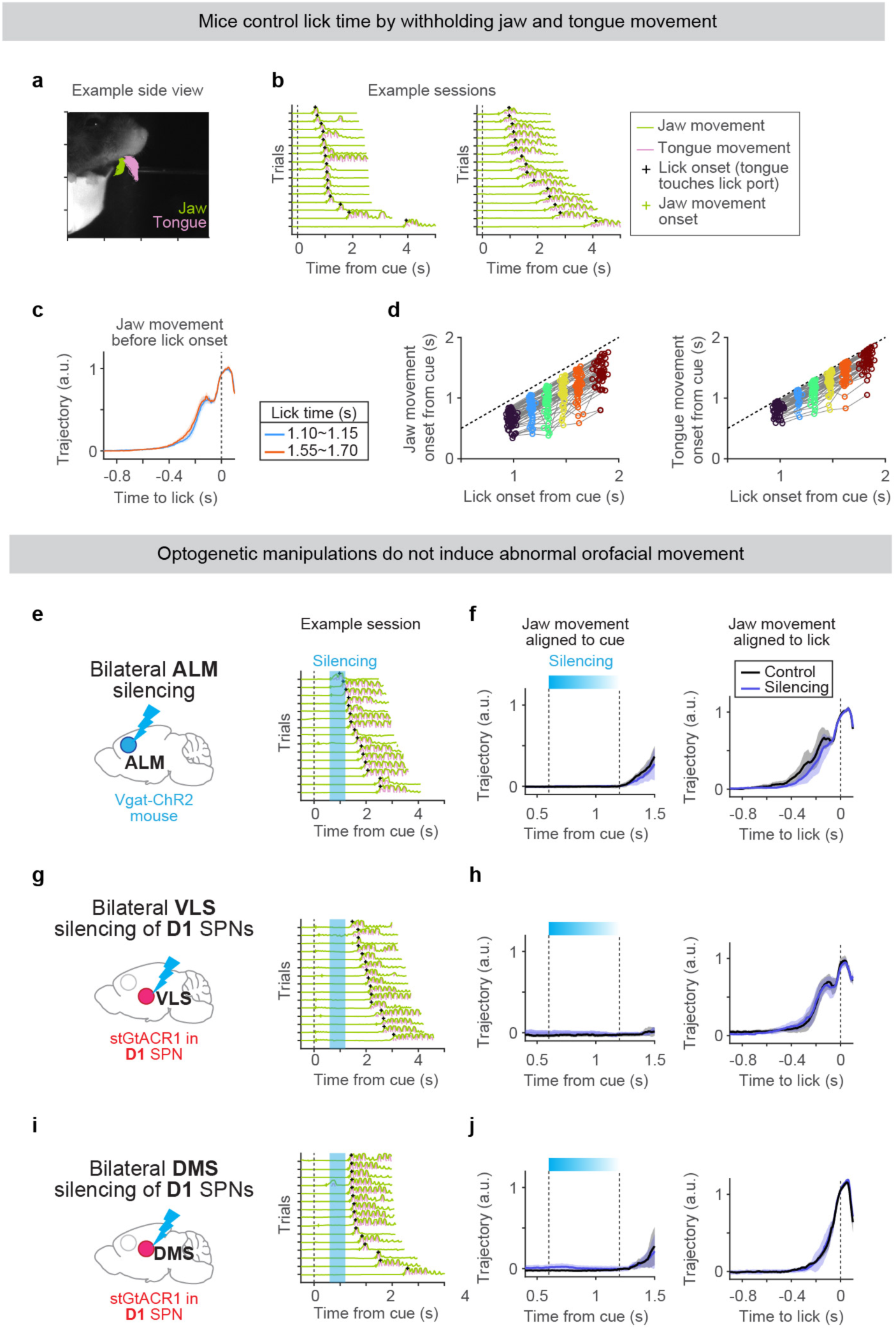
Characterization of orofacial movements in the lick-timing task. Jaw and tongue movements were tracked using high-speed videography and DeepLabCut^98^ (**a;** Methods). This revealed that mice adjusted lick times (detected by contact of the tongue with the lick port) by controlling the onset rather than the speed of tongue/jaw movements (**b-d**). Additionally, we did not detect any abnormal movements during/after ALM silencing (**e, f**), VLS inhibition (**g, h**), and DMS inhibition (**i, j**). **a.** An example side view clip of a mouse. Movement of the jaw (green) and tongue (purple) were tracked. Trajectories of individual trials are overlaid. **b.** vertical jaw (green) and tongue (purple) movements in two example sessions. The left session is from the animal shown in **a**. 15 randomly selected trials sorted by lick time are shown. **c.** AVerage trajectories of vertical jaw movement aligned to the lick onset. The kinematics of jaw movement remains consistent regardless of the lick timing (indicated in different colors). Lines, grand median. Shades, SEM (bootstrap). n = 58 sessions, 34 mice. **d.** Relationship between lick onset (the timing when the tongue contacted the lick port) and jaw movement onset (left) or tongue movement onset (right). Trials were grouped into six ranges. The onset of movement was tightly correlated with lick onset. Circles, individual mice. Dotted line, the unity line. N = 58 sessions, 34 mice. **c-d** conclude that mice did not change their kinematics but instead the onset of movement when they lick at different timings. n = 58 sessions, 34 mice. **e.** Tongue and jaw movement trajectories in an example session with ALM silencing. Same format as in **b**. Cyan bar, silencing. **f.** Jaw movement aligned to the cue (left) or the lick onset (right). Trials with lick after the silencing were analyzed. Lines, grand median. Shades, SEM (bootstrap). n = 26 sessions, 9 mice. No abnormal movement was detected during silencing, and the animals followed normal kinematics to lick even in the silencing trials. **g-h.** Same as in **e-f**, but with D1 VLS silencing. n = 6 sessions, 6 mice. **i-j.** Same as in **e-f**, but with D1 DMS silencing. n = 6 sessions, 6 mice.

**Extended Data Fig 4.**
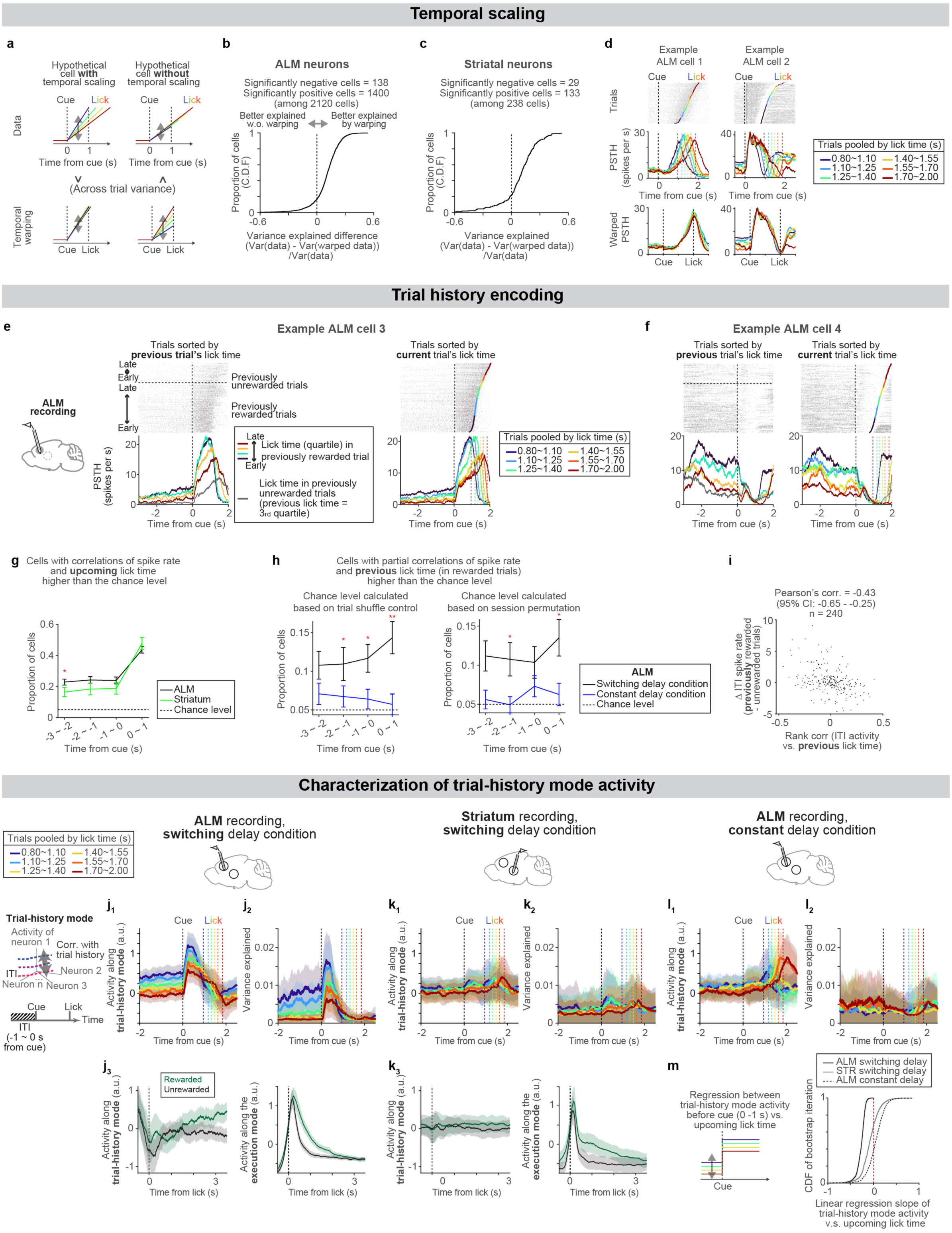
Temporal scaling and encoding of trial history in ALM and striatum. Temporal warping of spike times revealed that the activity in a large proportion of ALM and striatal neurons can be better explained by temporal scaling rather than a null model without scaling (**a-d**). Some ALM and striatal neurons maintain tonic activity during the ITI, which predicts upcoming lick time (**g**) and encodes trial history (**h-i** and example cells in **e** and **f**). Activity along the trial-history mode in the ALM under the switching delay condition predicts upcoming lick time (**j** and **m**). However, activity along the trial-history mode in the striatum under the switching delay condition (**k**) was weaker. Additionally, ALM activity under the constant delay condition (**l**) does not predict upcoming lick time, consistent with single cells (**h**). **a.** Schema illustrating two hypothetical cells that encode time differently through ramping activity^41^. Left, a cell with ramping activity encoding relative time, where the ramping speed changes as the lick time varies (i.e., temporal scaling). In this scenario, the across-trial variance (double-headed arrows) decreases following temporal warping (bottom). Right, a cell with ramping activity encoding absolute time, where the spike rate increases as time progresses. In this case, the across-trial variance increases following temporal warping (right). **b.** The cumulative distribution of the difference of variance explained between data and warped data across ALM neurons (higher value represents data explained better by temporal scaling; see **a**, Methods). n = 2139 neurons, 31 mice. Neurons with more than 200 trials were analyzed. Significant cells, p < 0.05 with bootstrap of trials. **c.** Same as in **b** but for striatal cells, n = 595 neurons, 10 mice. **d.** Two example ALM cells showing temporal scaling. Same format as Fig. 3a: trials are sorted by the current trial’s lick time and grouped into six ranges. **e.** An ALM example cell whose ITI activity is modulated by the lick time and reward outcome in the previous trial and anticipates upcoming lick time. The same cell as in Fig. 4a for comparison. Top, spike raster, grouped by reward outcome in the previous trial and sorted by the lick time in the previous trial. In this example cell, the ITI activity is higher in trials after rewarded trials and with earlier licks. Bottom, PSTH. Lick times of the previous rewarded trials were divided into quartiles indicated by different colors. The gray trace, trials following previously unrewarded trials with previous trial’s lick times within the 3rd quartile. Right, the same cell but trials sorted by lick time in current trials. The spike rate during ITI predicts the upcoming lick time. **f.** Another ALM example cell showing trial history modulation during ITI, same format as **e**. **g.** The proportion of neurons with a rank correlation between spiking activity (in different time windows indicated on the x-axis) and upcoming lick time higher than the trial shuffle control (*a* = 0.05). Neurons with more than 100 current trials and a current lick time later than 1 second (to avoid the influence of post-lick activity for the 0 - 1 second time window) were analyzed. * *p* < 0.05, hierarchical bootstrap comparing ALM and striatum. Bars, SEM (hierarchical bootstrap), **h.** The proportion of ALM neurons with a partial rank correlation between spiking activity (in different time windows indicated on the x-axis) and previous lick time higher than the trial shuffle control (left) and session permutation control (right) (*a* = 0.05). Partial correlation was calculated to control for the effect of upcoming lick time (Methods). Both yielded consistent results. Neurons with more than 50 current trials and a current lick time later than 1 second (to be consistent with g) were analyzed. * *p* < 0.05, ** p< 0.005, hierarchical bootstrap comparing switching vs. constant delay condition. Bars, SEM (hierarchical bootstrap), **i.** Relationship between the encoding of previous lick time (partial rank correlation between spike rate vs the lick time in previous rewarded trials, controlling for the effect of upcoming lick time; Methods) and whether the animal received a reward or not in the previous trial (based on activity during ITI: 0 - 1s before the cue) in ALM. Dots, individual neurons. Neurons with more than 20 previous unrewarded trials, 20 previous rewarded trials, 50 current trials with lick time later than 1 s were analyzed. Neurons encoding previous lick time also tend to encode previous reward outcome. **j.** ALM population activity along the trial-history mode, under switching delay condition (**j1**). Duplicated from Fig. 4b for comparison. variance of spiking activity explained by trial-history mode (**j2**). Colors, different lick times. Lines, grand mean. Shades, SEM (hierarchical bootstrap). n = 3261 neurons. ALM activity along the trial-history mode gradually diverged after rewarded vs. unrewarded licks (left) under the switching delay condition (**j3**). In contrast, ALM activity along the execution mode, which captures the activity during the lick (Methods), showed a transient change in activity after the lick (up to ∼2 s). This transient change along the execution mode most likely reflects differences in lick patterns between these trial types. Since the trial-history mode activity started diverging after the execution modes converged, the divergence of trial-history mode activity is likely not due to movement. Rewarded (green) and unrewarded (black) trials with similar lick times (lick between 1.4 and 1.8 s after the cue) were analyzed. **k.** Same as in **j** for striatal recording during the switching delay condition. **l.** Same as in **j** for ALM recording during the constant delay condition. **m.** The relationship between trial-history mode activity and upcoming lick time. We performed a linear regression analysis between trial-history mode activity during 0 - 1 s before the cue and the upcoming lick time for each iteration of the hierarchical bootstrap (left, schema). We then plotted the distribution of the linear regression coefficient (slope) across these iterations. A negative value indicates higher trial-history mode activity precedes earlier lick. *p* < 0.001, = 0.3274, 0.6310 for ALM under switching delay, striatum under switching delay, ALM under constant delay, respectively (with a null hypothesis that the slope of the linear regression is larger than or equal to 0). Thus, only under the switching delay condition in ALM, the trial-history mode activity significantly predicts the upcoming lick time.

**Extended Data Fig 5.**
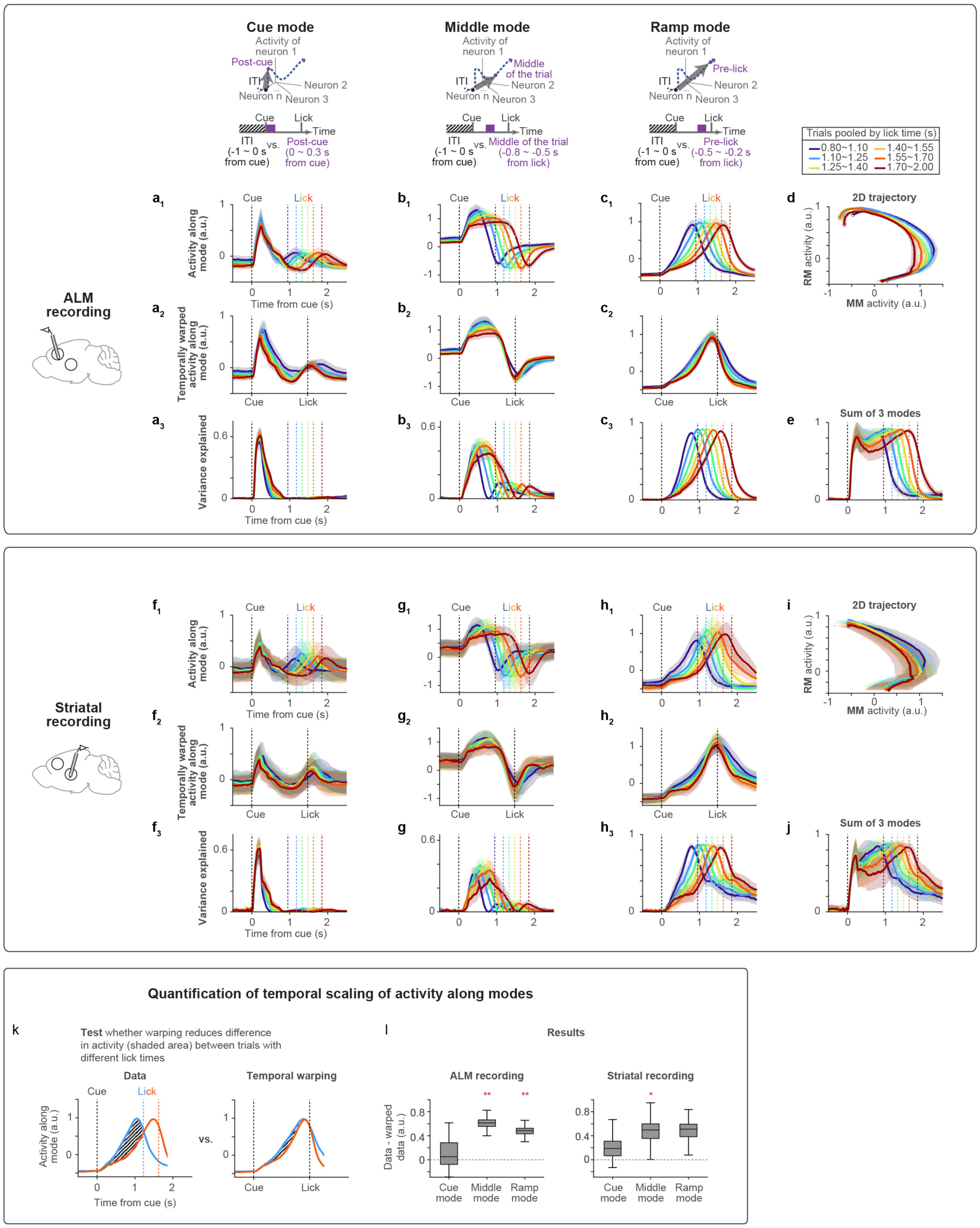
Comparison of activity modes across brain regions. In both ALM and striatum, we observed three modes of population activities (cue mode, middle mode, and ramp mode) that together tiled the time from trial onset to lick, explaining around 80% of the variance in task-modulated spiking activity. Activity along cue mode did not show temporal scaling, while activity along middle and ramp mode showed temporal scaling (quantified in **k** and **l**). Note that activity patterns along these modes and the variance explained are qualitatively similar between ALM and striatum. **a.** ALM population activity along the cue mode, under switching delay condition (**a1**). Cue mode activity temporally warped between cue and lick (**a2**, Methods). variance of spiking activity explained by cue mode (**a3**, Methods). Colors, different lick times. Lines, grand mean. Shades, SEM (hierarchical bootstrap). n = 3261 neurons. **b.** Same as in **a** but for ALM population activity along the middle mode. **c.** Same as in **a** but for ALM population activity along the ramp mode. Duplicated from Fig. 3e for comparison. **d.** Population activity in a two-dimensional space defined by the ramp mode (RM) and middle mode (MM). Trajectories are plotted from the cue (filled circles) to the lick (open circles). Regardless of the lick time, the activity follows a similar trajectory, but the speed varies across different lick times. **e.** The total variance explained by the three modes. **f-j.** Same as in **a-e** but for striatal neurons under switching delay condition. n = 1073 cells. **k.** Schema representing the quantification of temporal scaling. We calculated the difference in activity along a mode between two trial types (trials with licks occurring 1.1 - 1.25 seconds vs. 1.55 - 1.7 seconds; the difference was calculated from the cue to the lick, shaded area). If the population activity along a mode exhibits temporal scaling, the difference between lick times will be smaller following temporal warping. **l.** Left, the difference in activity (shaded area) as described in panel **k** is compared between the data and the temporally warped data for each mode in the ALM. Right, same for the striatum. The central line in the box plot, median. Top and bottom edges, 75% and 25% points. Whiskers, the lowest/highest datum within the 1.5 interquartile range of the lower/upper quartile. \*\**p* < 0.001, \**p* < 0.01 (significant scaling; hierarchical bootstrap with a null hypothesis that the difference between data minus temporal warped data is less than or equal to 0).

**Extended Data Fig 6.**
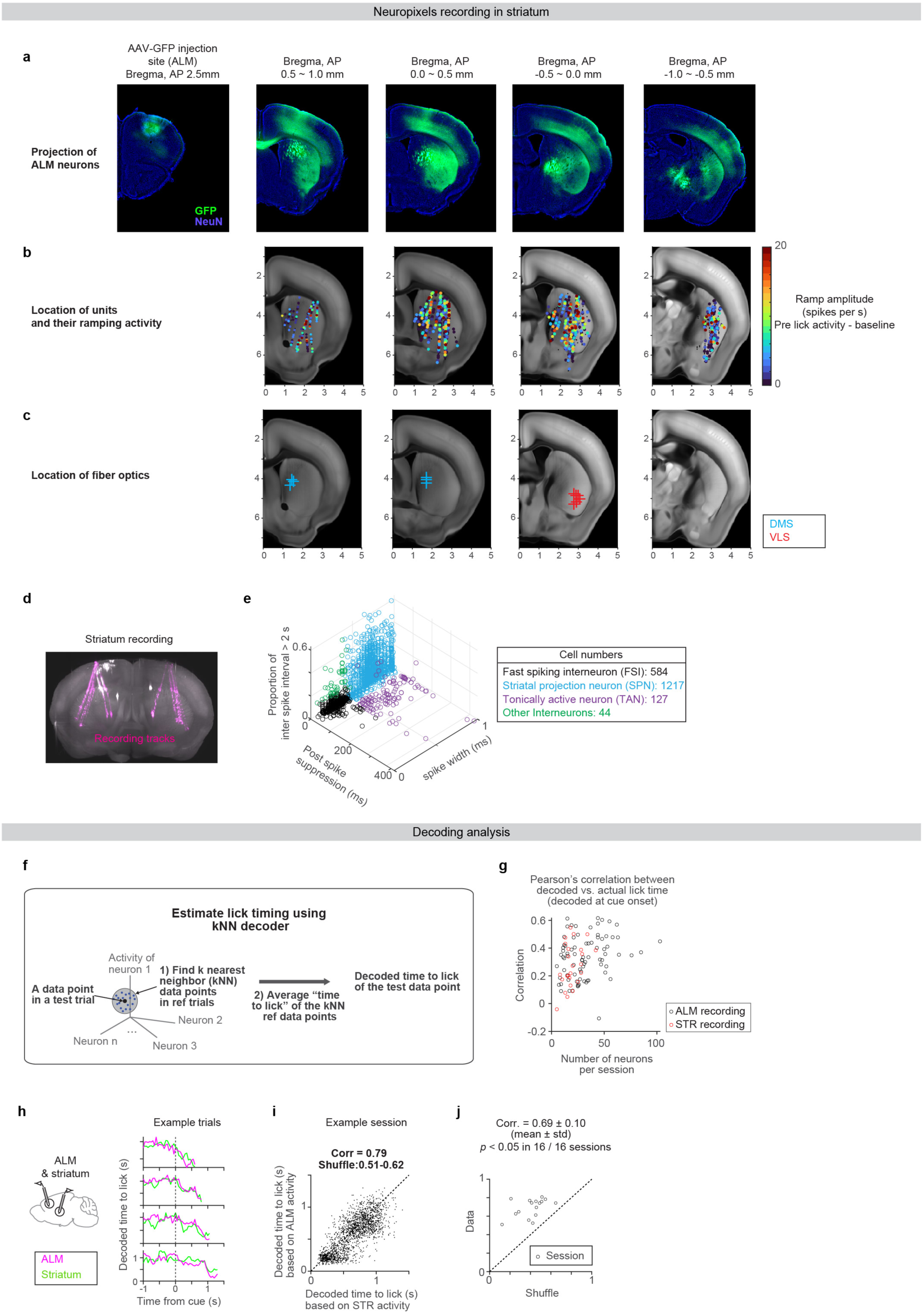
Striatal anatomy and recording. ALM projects to a large portion of the striatum (**a**), and we observed task-modulated activity across these sectors (**b**). See **d** for example recording tracks and **e** for the classification of cell types based on spike features. We developed a kNN-based decoder to estimate the time to lick using simultaneously recorded units at single time points (**f-g**). When ALM and striatum were simultaneously recorded, the decoded time to lick in both regions (decoded independently) was correlated at the single trial level, implying that the representations of time are highly synchronized between these areas (**h-j**). **a.** ALM neurons project across sectors in the striatum. AAV-GFP was injected into ALM. **b.** The spatial distribution of recorded striatal neurons in the Allen CCF. Colors, the extent of increase in spiking activity before the lick compared to the baseline. Black dots, neurons that do not ramp up. **c.** Locations of the tips of the tapered fiber optics implanted for bilateral striatal silencing (only showing one hemisphere as the two fibers were implanted symmetrically). Related to Fig.6. **d.** An example brain image of recording tracks acquired by a light sheet microscopy (Methods). Striatal recording tracks are labeled with CM-DiI (magenta). Coronal view, maximal intensity projection of 415 µm optical section. **e.** Striatal cell types were classified based on three spike features^57^ (Methods). **f.** Schema depicting a k-nearest neighbor (kNN) method to decode the time to lick (T_to lick_) using population neural activity at each time point (Methods). **g.** The performance of the kNN decoder as a function of the number of simultaneously recorded neurons. Decoding accuracy was quantified by Pearson’s correlation between actual lick time vs. lick time decoded at cue onset. The performance increased with more recorded neurons. **h.** Decoded T_to lick_ was highly correlated between ALM and striatum at a single trial level. Four example trials from an example session are shown. Traces end at the time of lick. **i.** The relationship between decoded lick times estimated from ALM and striatal neurons in an example session. Dots, all time points (50 ms bin; from cue to lick) in the example session. Pearson’s correlation across all time points (0.79) was significantly higher than that of the trial shuffle control (0.51 - 0.62, 95% confidence interval). **j.** Pearson’s correlation of decoded time between ALM and striatum was significantly higher than trial shuffle controls in all simultaneously recorded sessions (16 sessions). Thus, ALM and striatal timing dynamics are synchronized at a single trial level.

**Extended Data Fig 7.**
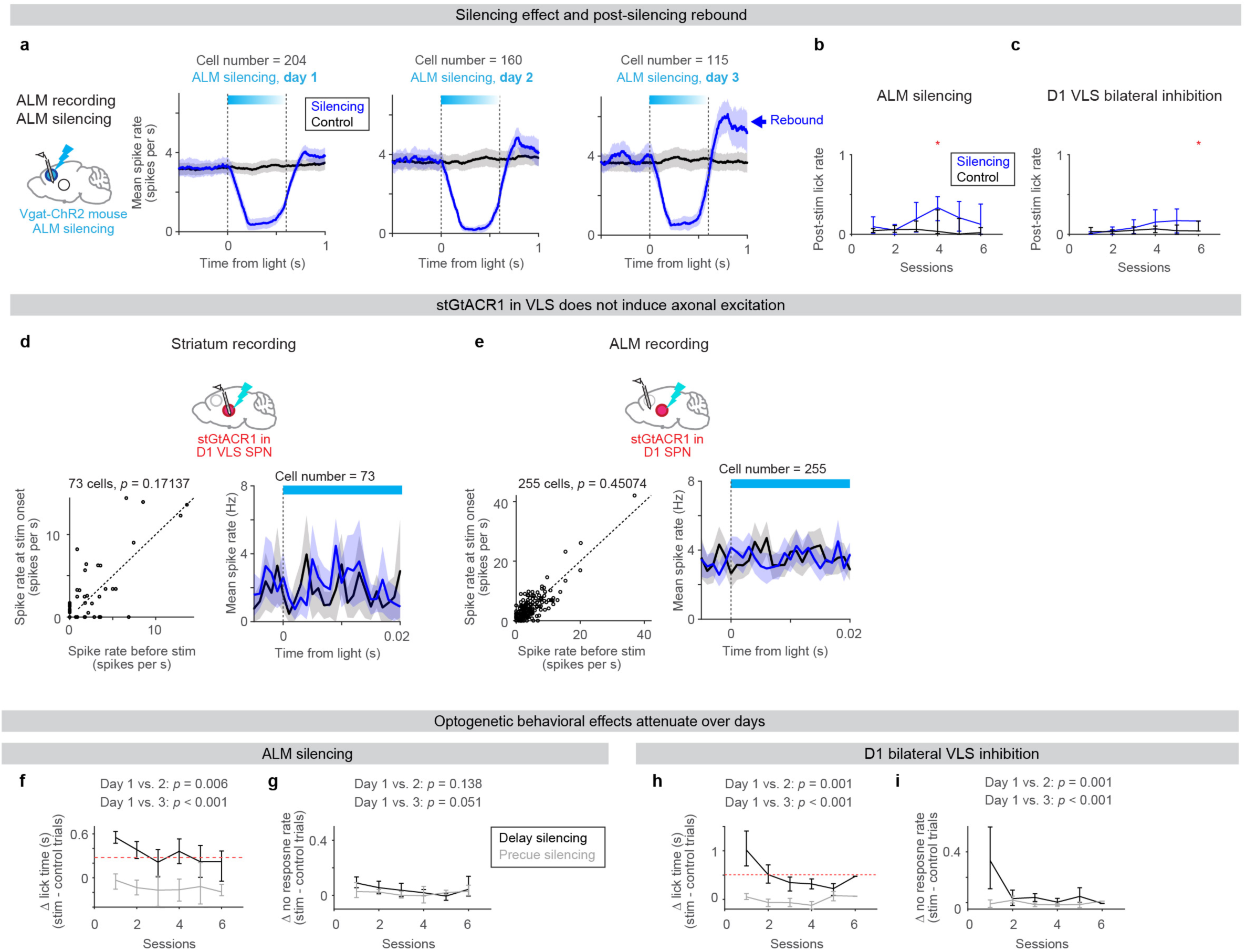
Characterization of optogenetic effect across sessions. Here, we characterized possible caveats of optogenetic manipulations. First, we measured the post-silencing rebound activity and lick, over days (**a-c**). We noticed that for ALM silencing, post-silencing rebound activity increased over days (**a**), accompanied by the increase of post-silencing lick (**b**), which explains the decrease in the shift of lick timing over sessions (**f-g**). Therefore, we restricted our analysis to data from the first two days. Second, we confirmed the lack of axonal excitation in both striatum and ALM when we used stGtACRl^59,76,77^ in D1-SPN (**d-e**). Third, we observed that the behavioral effect of D1-SPN inhibition showed a drastic decline over sessions, so we focused our analysis on the data from the first manipulation day (**h-i**). These observations are consistent with the short-lived effect of manipulations observed in other species and manipulations^61,62^. **a.** Mean spike rate of ALM putative pyramidal neurons during ALM silencing (during no cue trials) over 3 consecutive days (spike rate was smoothed by 200 ms boxcar causal filter). Lines, grand mean. Shades, SEM (hierarchical bootstrap). Note the increase in the post-silencing rebound activity over days (blue arrow). **b.** Post-stim lick rate (Proportion of trials with lick within 600 ms following the photostimulation offset time in no cue trials) across days for ALM silencing. \**p* < 0.001 (hierarchical bootstrap with *Bonferroni* correction for multiple comparisons; null hypothesis is that the post-stim lick rate in photostimulation trials is lower than or equal to that in control trials). **c.** Same as in **b** but for D1 VLS bilateral inhibition. **d.** When stGtACR1 exhibits leaky expression at the axon, it can induce short-latency axonal excitation at the onset of photostimulation^59,76,77^. Thus, we examined the spiking activity at the onset of D1-SPN inhibition (during no cue/precue silencing trials) in VLS, which did not show signs of axonal excitation. Left, spike rate during baseline (20 ms before the photostimulation) vs. spike rate within 20 ms of the photostimulation onset for individual striatal neurons. *P*-value, signed rank test. Right, mean spike rate of SPNs (1 ms bin, no smoothing of spike rate). n = 73 neurons. **e.** Same as in **d** but for ALM neurons during bilateral D1 VLS silencing. n = 255 neurons. Neither the striatum (**d**) nor ALM (**e**) showed signs of axonal excitation caused by GtACR1. **f.** The shift in lick time caused by ALM silencing became weaker over consecutive days with ALM silencing. Error bars, 95% confidence interval (hierarchical bootstrap). The red dashed line, half of the effect observed in session 1. *P*-value, bootstrap with a null hypothesis that the effect of delay silencing on session 1 is smaller than or equal to the subsequent compared day. n = 14 mice. **g.** The change in no response over consecutive days with ALM silencing. Error bar, 95% confidence interval (hierarchical bootstrap). *P*-value, bootstrap with a null hypothesis that the effect of delay silencing on session 1 is smaller than or equal to the subsequent compared day. **h-i.** Same as in **f**-**g** but for bilateral D1 SPN inhibition in VLS. n = 6 mice.

**Extended Data Fig 8.**
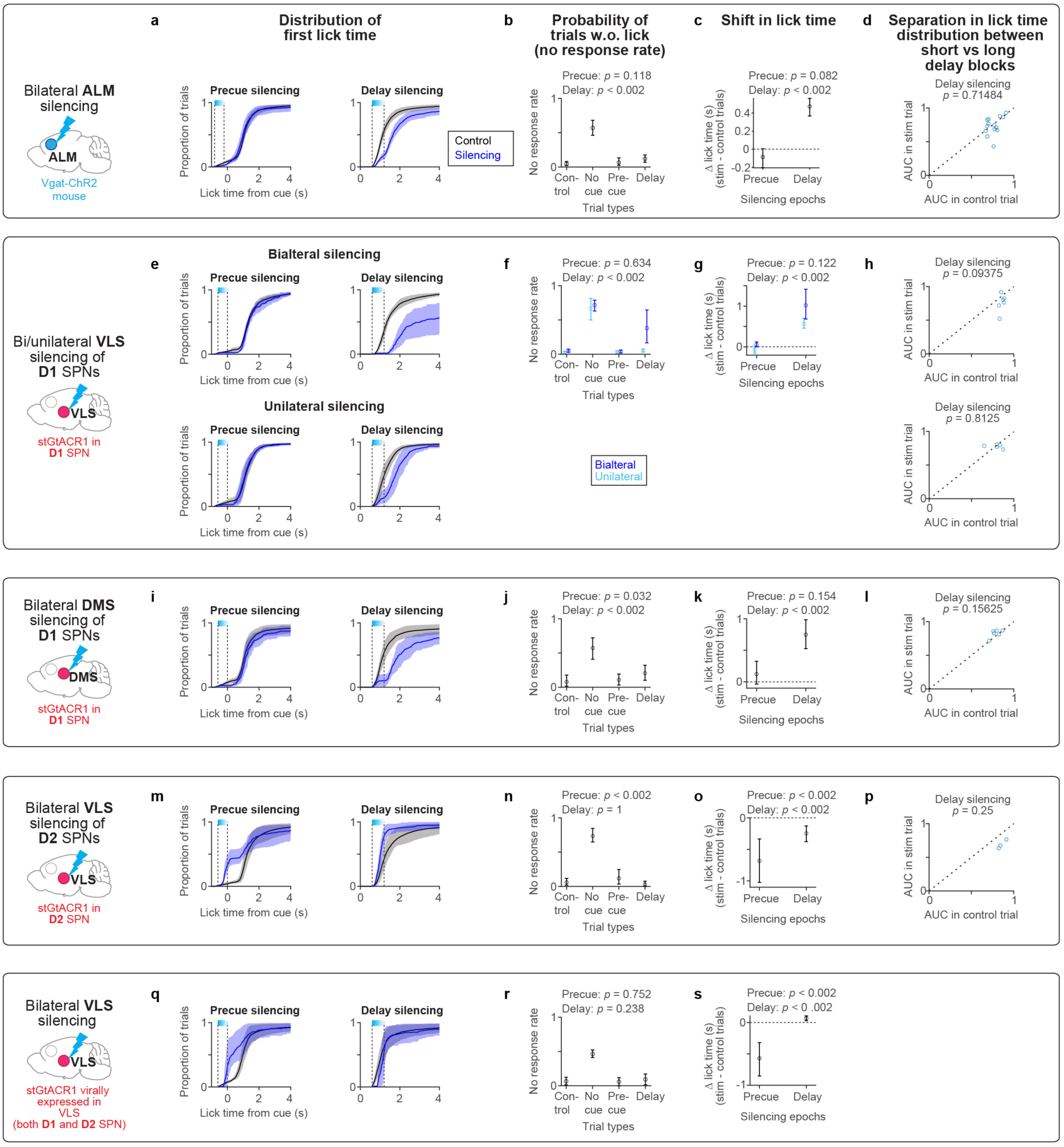
Summary of the behavioral effects observed across optogenetic manipulations. **a.** Lick time distribution during precue or delay ALM silencing, duplicated from Figure 5a for comparison. n = 14 mice. **b.** No-response rate in control, no cue, precue silencing, and delay silencing trials (Methods). No cue trial refers to randomly interleaved trials without a cue, serving to monitor the spontaneous lick rate not triggered by a cue. This represents the upper bound of the no-response rate. *P*-value, hierarchical bootstrap with a null hypothesis that no-response rate in control trials is the same as in silencing trials. Error bars, 95% confidence interval. **c.** Shift in median lick time caused by precue or delay silencing. Duplicated from Figure 5b for comparison. **d.** ALM silencing does not affect the separation in lick time distribution between delay blocks (short vs. long delay blocks in the switching delay condition). First, we calculated the area under the curve (AUC) of the ROC analysis comparing lick time distributions between short vs. long delay blocks. Then, we compared the AUC values (indicating how well the lick time distributions between delay blocks are separated) between control and photostimulation trials. Circles, individual animals. *P* value, signed rank test. The absence of change in the separation of lick time distributions suggests that ALM silencing does not erase the information of intended lick time, consistent with the recovery of dynamics after the silencing (Fig. 5). **e-h.**Same as in **a-d** but for D1 SPN inhibition in VLS. Top, bilateral D1 SPN inhibition in VLS (n = 6 mice). Duplicated from Figure 6 for comparison. Bottom, unilateral inhibition (n = 5 mice). **i-l.** Same as in **a-d** for D1 SPN inhibition in DMS. n = 6 mice. **m-p.** Same as in **a-d** for D2 SPN inhibition in VLS. n = 3 mice. **q-s.** Same as in **a-c** for cell-type-nonspecific striatal silencing in VLS. n = 2 mice.

**Extended Data Fig 9.**
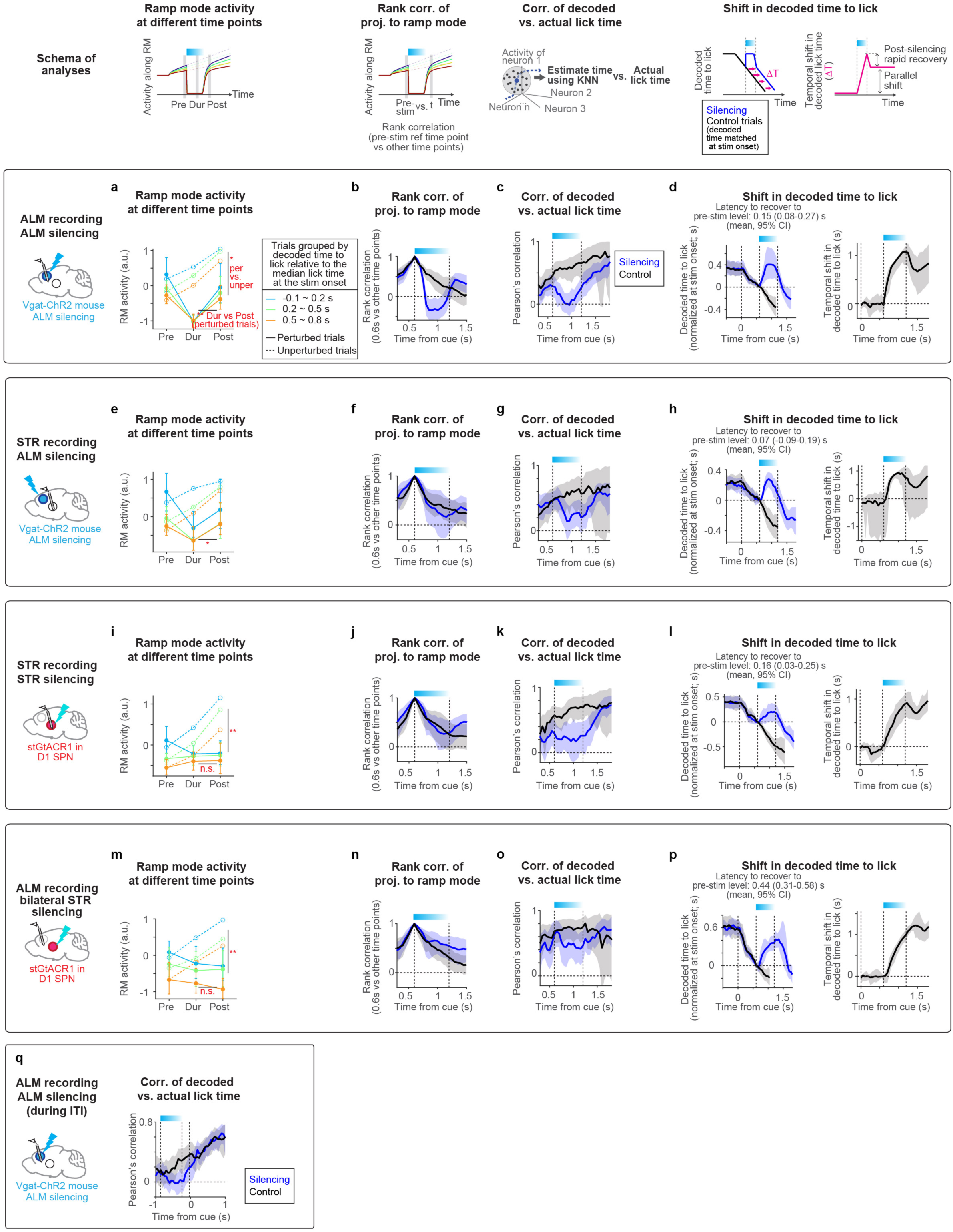
Quantification of timing dynamics across perturbation experiments. In addition to panels shown in Fig. 5 and 6, here, we present additional quantifications of the timing dynamics across perturbation experiments. Firstly, we quantified the change in RM activity during and after transient perturbation. Upon ALM perturbation, the RM activity in both ALM and striatum showed a V-shaped profile, reflecting rapid silencing and rapid recovery of the activity (**a** and **e**). This contrasts with the linear decay profile observed in both areas upon D1-SPN inhibition (**i** and **m**). Secondly, the rank correlation of projection along the ramp mode collapsed in ALM during ALM silencing (**b**), but not in other conditions (**f, j, and n**). Thirdly, the decodability of lick time (based on kNN decoder) collapsed in ALM during ALM silencing (**c**), but not in other conditions (**g, k, and o**). Fourthly, the decoded time to lick was shifted during perturbation and the shift persisted after the perturbation across conditions (**d, h, l, and p**). In contrast to silencing during the delay epoch, silencing before the cue was followed by a recovery of decoded lick time (**q**). **a.** Quantification of the change in RM activity during and after ALM silencing. RM activity at before (0.6 s after the cue, ‘Pre’), during (0.9 s after the cue, ‘Dur’), and after (1.25 s after the cue, ‘Post’) silencing is shown. Perturbed and unperturbed trials with matched T_to lick_ at the silencing onset (0.6 s after the cue) are shown. Different colors, trials with different decoded T_to lick_ relative to the median lick time in each session (to normalize for differences in median lick time across sessions). Dotted lines, unperturbed trials. Solid lines, perturbed trials. *P*-values, hierarchical bootstrap. **, *p* < 0.005; *, *p* < 0.05; n.s., non-significant. Circles, grand mean. Error bars, 95% confidence interval (hierarchical bootstrap). **b.** Rank correlation of ALM activity along the ramp mode across trials. The rank order at the pre-silencing condition (0.6 s after the cue) is compared with that at other time points. Lines, grand mean. Shades, 95% confidence interval (hierarchical bootstrap). The rank correlation in silenced trials (blue) collapsed during the silencing but recovered to the control (black) level afterward. **c.** Pearson’s correlation of lick time decoded by kNN decoder (based on ALM population activity) vs. the actual lick time. Lines, grand mean. Shades, 95% confidence interval (hierarchical bootstrap). ALM activity predicted upcoming lick time before and after the silencing, but such correlation disappeared during the silencing. **b** and **c** imply that after ALM silencing, the time information recovered. **d.** Decoded T_to lick_ from each time point based on kNN decoding analysis of population activity (left). Lines, grand mean. Shades, 95% confidence interval (hierarchical bootstrap). The decoded T_to lick_ was normalized by subtracting the decoded T_to lick_ at stimulus onset to account for different lick times across trials and sessions. Across conditions, the decoded T_to lick_ shifted during the perturbation, and the shift persisted after the perturbation, implying that the decoded T_to lick_ is shifted in parallel between control and silencing trials. **e-h.**Same as in **a-d** but for striatum recording during ALM silencing. **i-l.** Same as in **a-d** but for striatum recording during unilateral D1 SPN inhibition in VLS. **m-p.** Same as in **a-d** but for ALM recording during bilateral D1 SPN inhibition in VLS. **q.** Same as in **c** but for ALM recording during ALM precue silencing. Note that the correlation decays during the silencing, but recovered after the silencing.

**Extended Data Fig 10.**
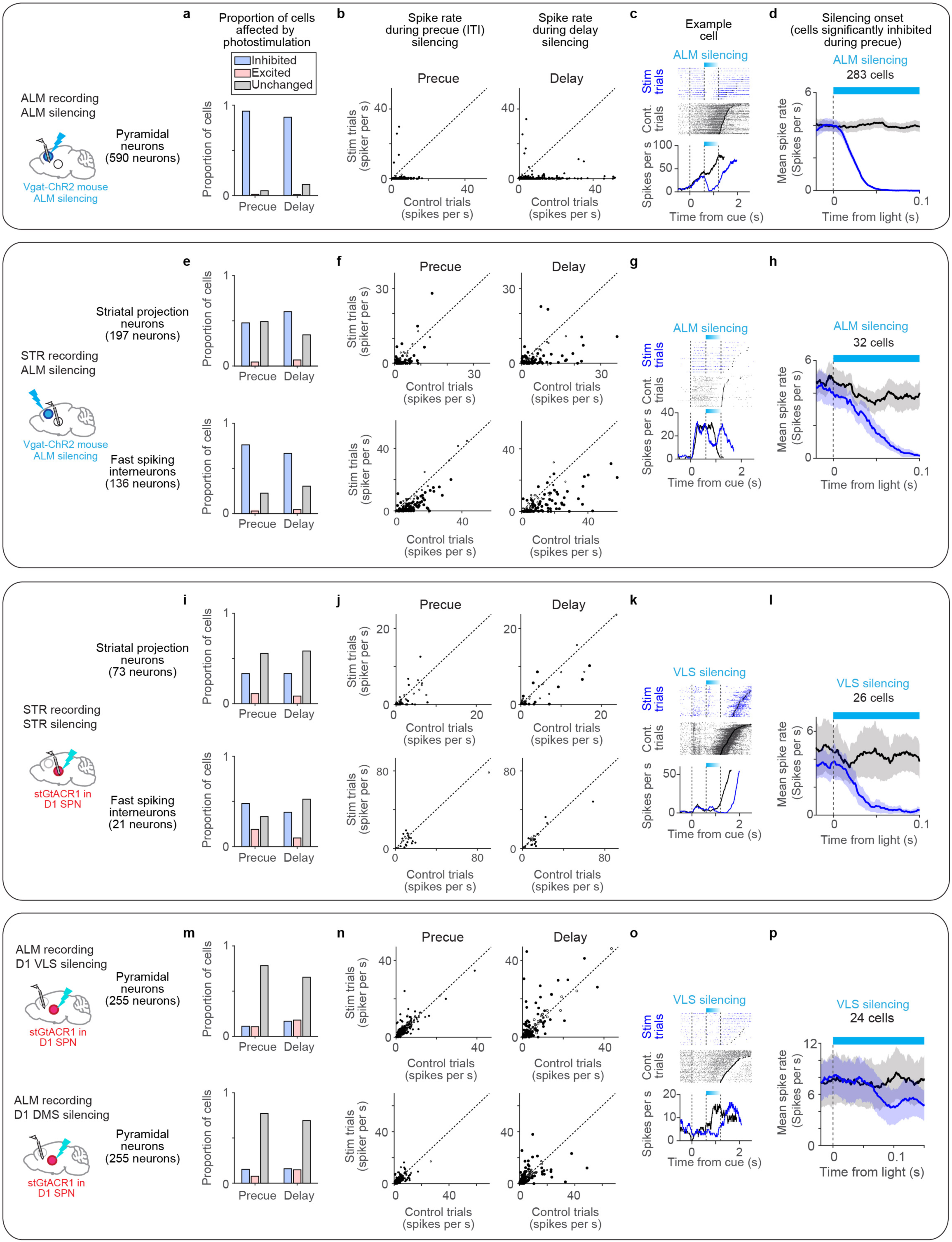
Summary of optogenetic effect on spiking activity. We characterized the effects of different optogenetic manipulations on spiking activity, including the proportion of inhibited/excited/unaffected cells (**a, e, i,** and **m**), changes in spike rate (**b, f, j,** and **n**), and the onset of the optogenetic effect in significantly inhibited neurons (analyzed during the precue epoch to avoid confounds caused by behavior-related activity; **d, h, l,** and **p**). **a.** The proportion of ALM pyramidal neurons affected by bilateral ALM silencing was assessed. Neurons were categorized as inhibited or excited (*p* < 0.05) or unchanged (*p* > 0.05) using a rank sum test. Cells with a mean spike rate higher than 1 Hz in control trials during the photostimulation window were considered. Duplicated from Figure 5c for comparison. Units from the first two manipulation sessions were included. **b.** Spike rate of ALM neurons during precue (left) or delay silencing (right) compared to control unperturbed trials. Circles, cells. Filled circles, significantly affected cells (*p* < 0.05, rank-sum test). The spike rate was calculated between 50 - 250 ms from the photostimulation onset. **c.** An example ALM neuron with a reduced spike rate during delay ALM silencing. Top, spike raster. Blue, silencing trials. Black, control trials. Bottom, PSTH up to the median lick time for each trial type. The spike rate was smoothed using a 200 ms boxcar causal filter. **d.** The onset of the optogenetic effect in ALM neurons with a significant reduction in spike rate during ALM silencing (*p* < 0.05, rank sum test). We analyzed silencing during the precue epoch to avoid confounds caused by behavior-related activity. Neurons with more than 10 trials and a spike rate higher than 1 Hz in the control condition were considered. The spike rate was smoothed using a 30 ms boxcar causal filter and aligned to the photostimulation onset. Blue, silencing trials. Black, control trials. Lines, grand mean. Shades, SEM (hierarchical bootstrap). **e-h.**Same as in **a-d** but for striatal recording during ALM silencing. Top, striatal projection neurons. Bottom, striatal fast-spiking interneurons. Units from the first two manipulation sessions were included. **i-l.** Same as in **a-d** but for striatal recording during D1 SPN unilateral inhibition. Top, striatal projection neurons. Bottom, striatal fast-spiking interneurons. Data from the first session was included for **i-k**. However, for **l**, data was pooled across 3 sessions, as the onset of silencing did not appear to change across days, despite significant behavioral changes likely caused by rebound or other activity changes after the silencing. **m-p.** Same as in **a-d** but for ALM recording during D1 silencing in VLS (top), or in DMS (bottom). Units from the first manipulation session were included.

**Extended Data Fig 11.**
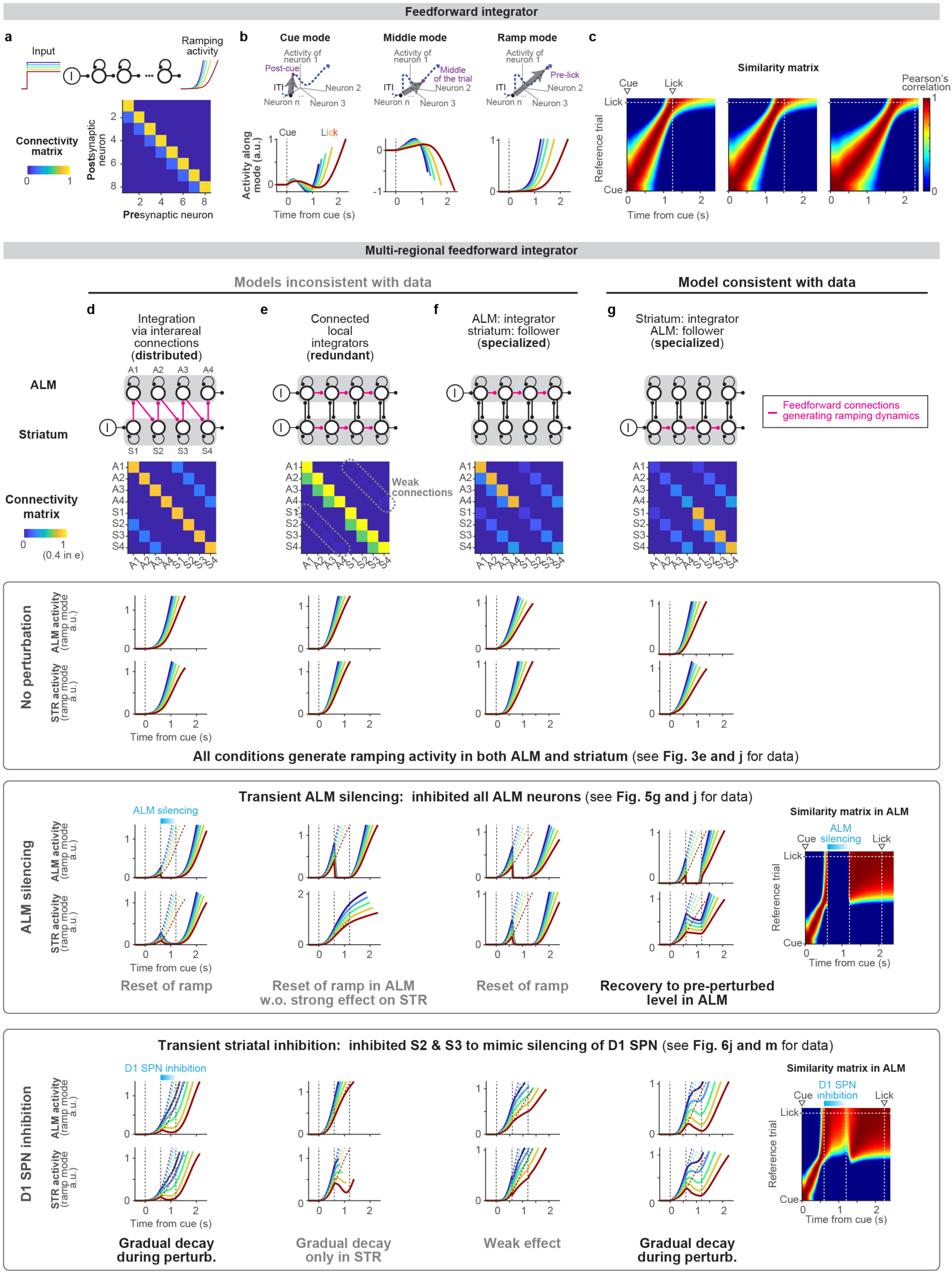
Multi-regional feedforward integrator. Recurrent networks with feedforward connections can temporally integrate step input to generate ramping activity^67^ (**a**; Methods), similar to an integrator based on positive feedback connections (Extended Data Fig. 1). Activity in this network is high-dimensional and exhibits gradual changes in population activity patterns over time akin to the data (**b-c**; see Methods). To analyze perturbation effects on the multiregional feedforward (FF) integrator models with distinct computational roles of ALM and striatum (STR), we placed FF connections differently (**d-g**). Consistent with the results in the positive feedback integrator (Extended Data Fig. 1), ALM functioning as an input/follower of the striatal integrator mimics the data, with a pause and rewind in the representation of time during ALM and striatal inhibition, respectively. **a.** Schema and connectivity matrix in the feedforward integrator network. **b.** Activity along different modes in the network described in **a**. **c.** Similarity matrix of population activity patterns in the network described in **a**. **d.** A multi-regional network where FF connections are distributed both in ALM and STR. The format is identical to that in Extended Data Fig. 1. In brief, to replicate ALM complete silencing, we injected a strong negative current into all ALM neurons (a1-a4). To replicate D1-SPN inhibition, we injected negative currents into half of the STR neurons (s2 and s3). Plots are shown up to the time of the lick (when ramping activity reaches a threshold level; the threshold was adjusted so that the lick occurs approximately 1 second after the cue in unperturbed trials). **e.** Similar to **c**, but a multi-regional network where two FF integrators in ALM and STR are weakly coupled. The perturbation of one area has only a weak effect on the representation of time in the other area. **f.** Similar to **c**, but a multi-regional network where FF connections reside in ALM, and STR provides feedback amplification of the signal. This allows STR to have similar dynamics as ALM, and ALM dynamics cannot progress without STR’s recurrent connection. STR feedback supplies drive for the ALM FF network to be functional. Perturbation of ALM resets the integration. **g.** Similar to **f**, but with an opposite configuration, i.e., FF connections in STR and ALM mediate feedback amplification of the signal. As ALM recurrent input supplies drive for the STR FF dynamics to progress, ALM silencing results in a pause in the representation of time. In contrast, STR inhibition (targeting half of STR neurons as in the data) rewinds the representation of time.

**Extended Data Fig 12.**
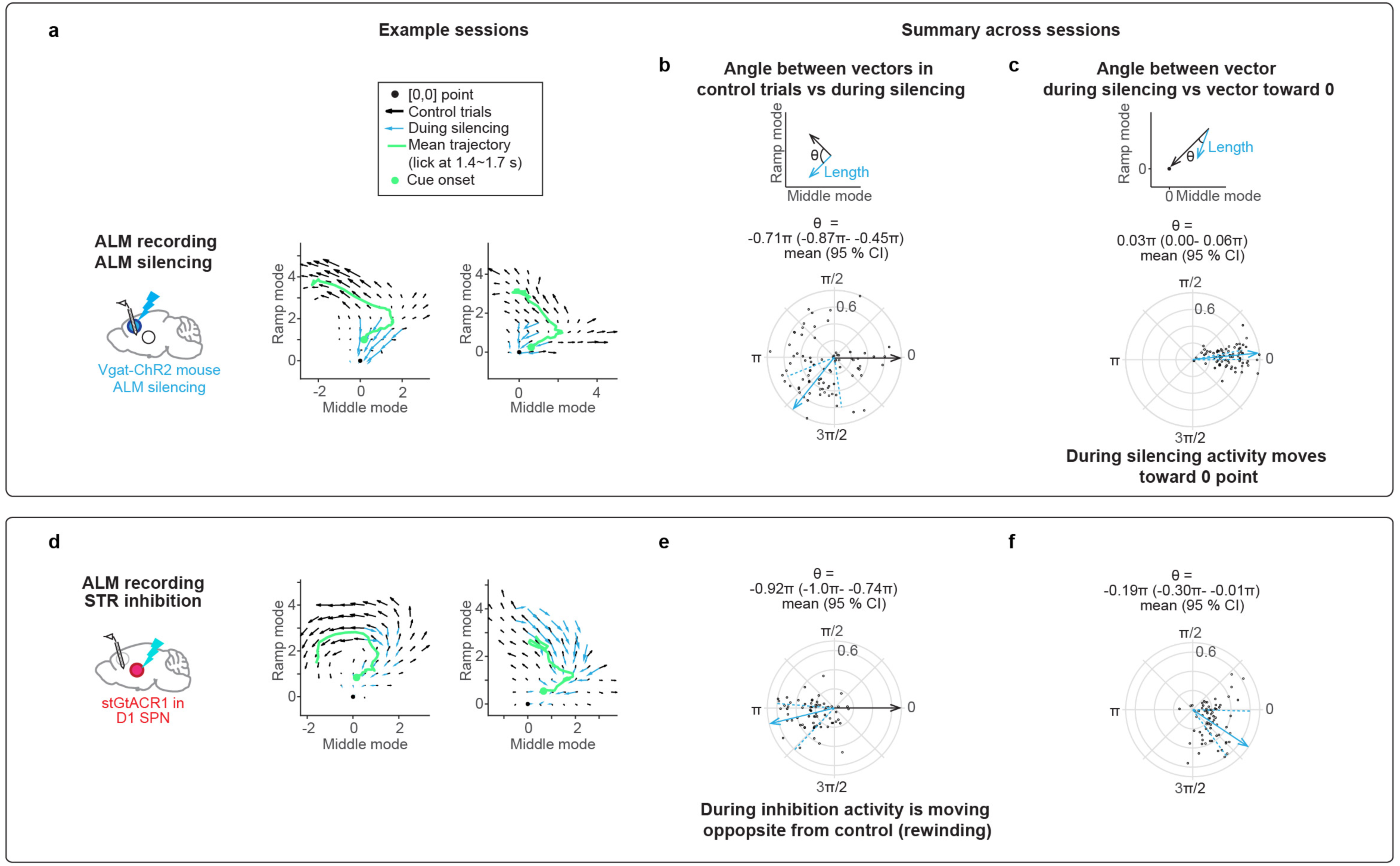
Rewinding of ALM dynamics during D1-SPN silencing in VLS. We analyzed how activity evolves in the two-dimensional activity space defined by RM and MM, which captures a large fraction of task-modulated activity (Extended Data Fig. 5). For each session, we calculated the direction in which the activity developed from individual activity states. Specifically, we pooled all activity states (50 ms bin) between cue and lick in unperturbed control trials, calculated the direction of activity evolution (in the next 50 ms bin), and averaged these to acquire the vector field per session (black arrows in **a** and **d**; for states with more than 30 data points). Similarly, we acquired the vector field during inhibition by pooling all activity states during inhibition in silenced trials (cyan arrows; Methods). We then calculated the angle between vectors in control versus inhibition (**b** and **e**; circles, individual states with both black and cyan arrows) to test whether activity evolved in the opposite direction from the normal trajectory during inhibition. Additionally, we calculated the angle between vectors during silencing and the vector toward the zero point (where the spike rate is 0) to test whether activity evolved toward the zero point (**c** and **f**). During ALM silencing, the trajectory is better explained by activity moving toward zero (compare **b** vs. **c**). In contrast, during D1-SPN silencing, the trajectory is better explained by activity moving in the opposite direction from the normal trajectory (compare **e** vs. **f**).

**Extended Data Fig 13.**
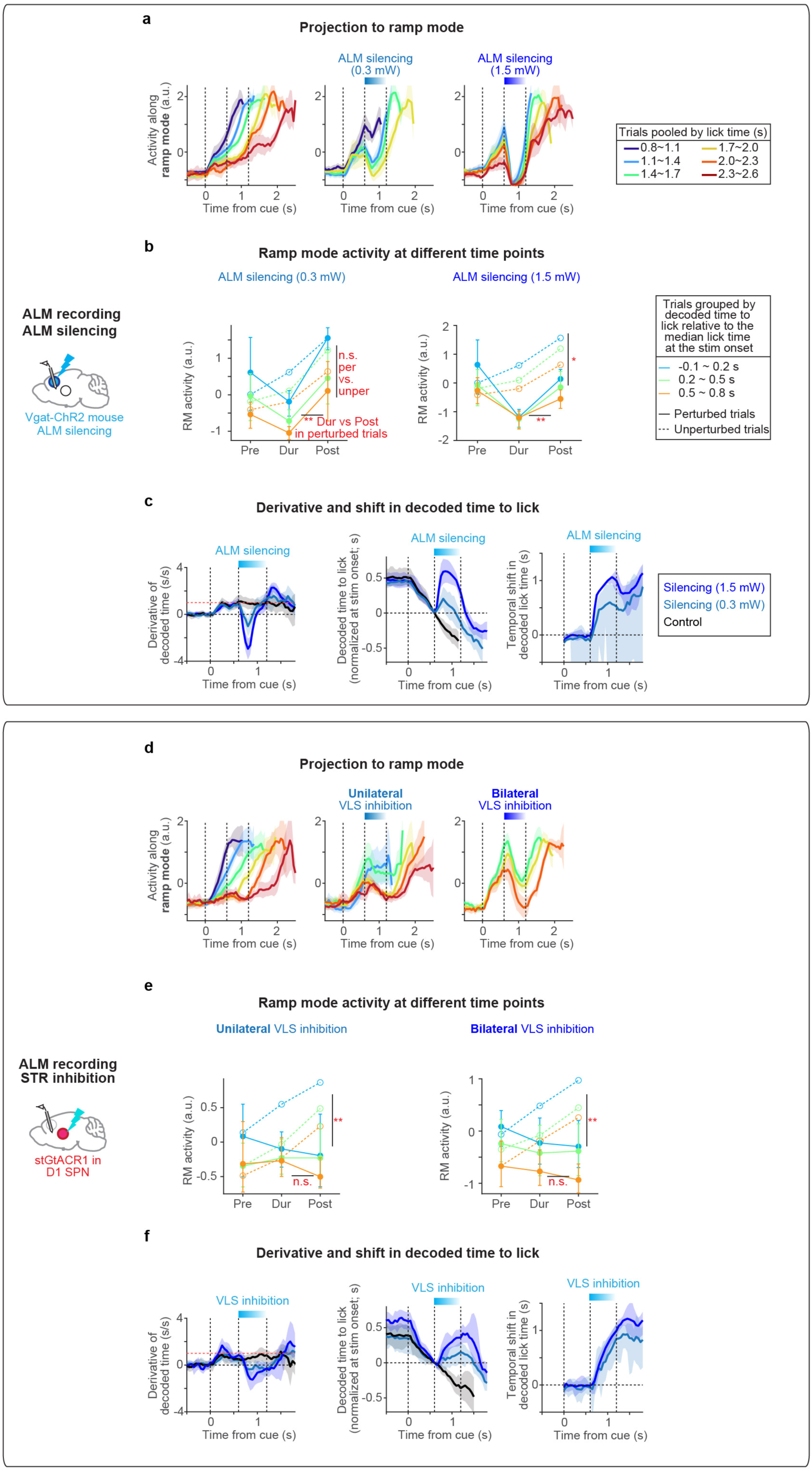
Graded and persistent effect of perturbations. We recorded ALM activity while perturbing it with different laser powers (0.3 vs 1.5 mW; **a-c**) or while unilaterally (ipsilateral) or bilaterally inhibiting D1-SPN in the VLS (**d-f**). In both cases, the perturbation with different intensities resulted in graded and long-lasting changes in ramping activity and decoded time beyond the duration of perturbations, consistent with the notion that the perturbation was integrated into the timing dynamics. **a.** ALM population activity along the ramp mode in control trials (left) and ALM silencing trials with different powers (middle, right). Lines, grand mean. Shades, 95% confidence interval (hierarchical bootstrap). n = 130 cells, 4 mice. Trials with different laser powers were randomly interleaved in the same sessions. **b.** Quantification of the change in RM activity during and after ALM silencing. Same format as in Extended Data Fig. 9a. Left, weak ALM silencing. Right, strong ALM silencing. Note that the plot of perturbed trials exhibited a V-shaped profile regardless of laser power, whereas in D1-SPN inhibition (**e**), it showed a linear decay profile regardless of whether the inhibition was unilateral or bilateral. **c.** Analysis of decoded time based on ALM population activity (using kNN decoder). Left: derivative of decoded time (computed over a 200 ms window). In control trials (block), the derivative changes from 0 to 1 (red horizontal dashed line) after the cue. In stimulated trials, the derivative becomes negative at light onset, reflecting rapid decay in activity, and increases after the light, reflecting rapid recovery. Middle and right: same format as in Extended Data Fig 9d. Lines, grand mean. Shades, 95% confidence interval. **d-f.** Same as in **a-c** for unilateral vs. bilateral inhibition of D1-SPNs in VLS. Data from different mice were combined (bilateral data is duplicated from Extended Data Fig. 9m; unilateral data, n = 113 cells, 4 mice). Unlike ALM silencing, the derivative of decoded time does not show a large change even during inhibition, reflecting a gradual change in timing dynamics.

