## Extended Data Table 1 for "Integrator dynamics in the cortico-basal ganglia loop underlie flexible motor timing"

| Experiment | Mice | Sessions | Total neurons | Putative pyramidal /striatal projection neurons | Corresponding figures |
|---|---|---|---|---|---|
| **ALM recording under switching delay condition** | | | | | |
| Bilateral ALM silencing | 14 (VGAT-ChR2-EYFP) | 28 | 711 | 590 | Fig. 5c and e-g, EDF. 7a, EDF. 9a-d,q, EDF. 10a-d, EDF. 12a-c, and EDF. 13a-c |
| Bilateral ALM weak silencing | 5 (VGAT-ChR2-EYFP) | 10 | 225 | 193 | EDF. 13a-c |
| Bilateral VLS D1 silencing | 6 (Drd1-cre FK150 x R26-LNL-GtACR1-Fred-Kv2.1) | 6 | 283 | 255 | Fig. 6e, and k-m, EDF. 7e, EDF. 9m-p, EDF. 10m-p, EDF. 12d-f, and EDF. 13d-f |
| Unilateral VLS D1 silencing | 4 (Drd1-cre FK150 x R26-LNL-GtACR1-Fred-Kv2.1) | 4 | 320 | 282 | EDF. 13d-f |
| Bilateral DMS D1 silencing | 6 (Drd1-cre FK150 x R26-LNL-GtACR1-Fred-Kv2.1) | 6 | 287 | 255 | EDF. 10 mn, bottom |
| Other recordings | 8 | 108 | 3267 | 2892 | N/A |
| The sum of all experiments | 43 | 162 | 5093 | 4467 | Fig. 3b-e, Fig. 4b, EDF. 4g-j and m, and EDF. 5a-e and l |
| **Striatal recording under switching delay condition** | | | | | |
| Bilateral ALM silencing | 7 (VGAT-ChR2-EYFP) | 13 | 372 | 197 | Fig. 5c, h-j, EDF. 9e-h, and EDF. 10e-h |
| Unilateral VLS D1 silencing | 5 (Drd1-cre FK150 x R26-LNL-GtACR1-Fred-Kv2.1) | 10 | 103 | 73 | Fig. 6e, h-j, EDF. 7d, EDF. 9i-l, and EDF. 10i-l |
| Other recordings | 4 (VGAT-ChR2-EYFP)+ all mice above | 74 | 1497 | 947 | N/A |
| The sum of all experiments | 16 | 97 | 1972 | 1217 | Fig. 3g-j, EDF. 4g, k and m, and EDF. 5f-j and l |
| **ALM recording under constant delay condition** | | | | | |
| The sum of all experiments | 13 (7 C57Bl/6J; 6 VGAT-ChR2-EYFP) | 63 | 2609 | 2280 | EDF. 4h, l and m |

**Extended Data Table. 1: sample sizes (n) in the paper**

Neurons used for analysis in the rows of “The sum of all experiments” were based on whether the neuron has more than 10 trials for all 6 lick time ranges: 0.80~1.10, 1.10~1.25, 1.25~1.40, 1.40~1.55, 1.55~1.70, 1.70~2.00 sec.
